# Forward-projected cortical eigenmodes provide an efficient sensor-space representation of resting-state EEG

**DOI:** 10.64898/2025.12.08.693061

**Authors:** Hyung G. Park

## Abstract

Sensor-space EEG analyses typically rely on electrode layouts or data-driven components and rarely encode cortical geometry, making scalp patterns difficult to link to anatomy and to compare across participants. We introduce a sensor-space basis dictionary that explicitly integrates cortical geometry. Laplace–Beltrami (LB) eigenmodes are computed on a standard cortical template (fsaverage) and mapped by the lead-field matrix of a three-layer boundary-element (BEM) head model to yield cortex-anchored sensor-space harmonics. The leadfield-mapped LB dictionary spans scalp topographies, while preserving a meaningful spatial-frequency ordering inherited from the cortical manifold. We assess representational efficiency using ordinary least squares (OLS) projections of resting EEG (eyes-closed/open) across 59-, 32-, and 19-channel montages, and compare against spherical harmonics (SPH), principal components (PCA), and independent components (ICA). Efficiency is quantified by the variance explained *R*^2^(*K*) (by leading *K* modes) and the efficiency indices *K*_70_ and *K*_90_ (fewest modes reaching *R*^2^ ≥ 0.70 and 0.90) and reliability by ICC(3,1) of eyes-open/closed coefficients. The cortex-anchored basis shows higher early-*K R*^2^ than SPH and PCA (e.g., 59-channel eyes-closed at *K*=4: LB *R*^2^ ≈ 0.56 [95% CI: 0.54, 0.59] vs. SPH ≈ 0.44 [0.42, 0.46], PCA ≈ 0.08 [0.07, 0.09]) and reaches 70% and 90% variance with fewer modes (LB *K*_70_ ≈ 8.6; SPH ≈ 12.3; PCA ≈ 19.3; ICA ≈ 22.8; LB *K*_90_ ≈ 22.6; SPH ≈ 25.3; PCA ≈ 23.2; ICA ≈ 30.2). Mode-wise coefficient reliability (eyes-open vs. eyes-closed) is comparable between LB and SPH. By combining cortical eigenmodes with a forward head model, this approach yields a geometry-aligned, interpretable representation of sensor-space EEG that offers superior fidelity-complexity trade-offs at small *K* and a principled scaffold for low-dimensional EEG sensor space analysis.

## Introduction

Electroencephalography (EEG) is inexpensive, portable, and temporally precise, but its spatial specificity is fundamentally limited by volume conduction and the ill-posedness of the inverse problem: many distinct cortical current configurations produce nearly indistinguishable scalp fields (Nunez and Srinivasan, 2006; Hämäläinen et al., 1993). The poorly conducting skull acts as a strong low-pass spatial filter, so fine-scale cortical detail is blurred at the sensors and a large fraction of resolvable variance resides at coarse spatial scales (Srinivasan et al., 1998; Dannhauer et al., 2011; Iivanainen et al., 2021b). These limits motivate analyzing the scalp electric field as a spatial field—rather than as single channels—to recover interpretable structure (Michel and Murray, 2012). A principled alternative is to constrain analysis to a *structured* set of functions that (i) lies within the subspace supported by the head-tissue volume conductor and (ii) orders components from coarse to fine spatial scale. Data-adaptive decompositions such as principal component analysis (PCA) (Jolliffe, 2002; Dien, 2012) and independent component analysis (ICA) (Hyvärinen, 1999; Hyvärinen and Oja, 2000; Delorme and Makeig, 2004) are powerful but not inherently anatomical without additional constraints (Haufe et al., 2014; Sassenhagen and Draschkow, 2019), reinforcing the need for geometry-aware priors in sensor space.

Classic sensor-space parameterizations, such as spherical harmonics, are useful for interpolation and for constructing geometry-agnostic scalp bases (Perrin et al., 1989); however, they ignore the folded cortical geometry that generates EEG and the mixing imposed by the lead field. In contrast, *cortical* Laplace–Beltrami (LB) eigenmodes encode spatial scale directly on the *cortical surface* and have been used, together with related graph/spectral constructions, to explain large-scale spatiotemporal patterns across imaging modalities (Pang et al., 2023; Glomb et al., 2020; Henderson et al., 2022; Atasoy et al., 2016; Robinson et al., 2016). A small number of low-order modes often accounts for a substantial share of variance, reflecting the link between geometry and function and the role of cortical geometry as a scaffold for macroscopic neural dynamics.

### From cortex to sensors

We bring this idea directly into EEG sensor space. Starting from Laplace–Beltrami (LB) eigenmodes on the FreeSurfer *fsaverage* cortical surface, we map each *cortical harmonic* through a realistic three-layer lead-field model to obtain a *cortex-anchored* sensor-space basis: the columns of our dictionary are the scalp topographies generated by individual cortical harmonics. Unlike spherical harmonics defined purely in sensor geometry, this construction respects both the folded cortical anatomy and the physics of volume conduction, while remaining entirely in sensor space (no subject-specific inverse modeling required). Our approach retains the anatomical scaffolding championed in EEG imaging work while remaining purely in sensor space (Michel and Murray, 2012).

### Goal and contribution

We introduce and evaluate a *sensor-space LB dictionary* : LB eigenmodes computed once on a standard cortical template (FreeSurfer *fsaverage*) and mapped to the electrodes via a three-layer boundary-element (BEM) leadfield (Fischl, 2012; Fuchs et al., 2002; Mosher et al., 1999). On resting-state EEG (eyes-closed/open) and commonly used 19-, 32-, and 59-channel 10–20/10–10 montages (Klem et al., 1999; Oostenveld and Praamstra, 2001), we test whether this geometry-aligned basis (i) achieves higher representational efficiency, in terms of group-mean *R*^2^(*K*) as a function of the number of modes *K* and the efficiency indices *K*_70_ and *K*_90_ (fewest modes with *R*^2^ ≥ 0.70 and 0.90), in comparison to spherical harmonics (SPH) and data-adaptive PCA/ICA, and (ii) yields reliable coefficient structure across conditions (EO vs. EC), indexed by ICC(3,1) (Shrout and Fleiss, 1979). Observed time–frequency scalp maps are projected by ordinary least squares onto an orthonormal basis spanning the LB-induced sensor subspace, enabling rank-aware, montage-matched representations without subject-specific MRIs or source reconstructions. Our results support forward-projected cortical eigenmodes as a cortical geometry-aligned parsimonious coordinate system for sensor-space EEG.

## Methods and Experimental Plan

### Dataset

We analyzed resting-state EEG from the **Leipzig Mind–Brain–Body (MPI–LEMON)** dataset (Babayan and et al., 2019). EEG was recorded with a 62-channel EasyCap montage (61 scalp EEG plus one vertical EOG) at 1 kHz, with separate eyes-closed (EC) and eyes-open (EO) runs for each participant. Structural MRI (T1w) is available in MPI– LEMON but was not used to form subject-specific bases. We used both EC and EO to quantify (i) representational efficiency as a function of the number of basis modes and (ii)across-condition reliability (EO vs. EC) of mode coefficients.

#### Preprocessing and montage harmonization

We used MPI–LEMON’s publicly released, BIDS-organized (Pernet et al., 2019) *preprocessed* EEG (downsampling, band-pass filtering, channel cleaning, ICA-based ocular/ECG removal) without additional artifact correction. *Boundary-aware windowing:* EC and EO recordings are concatenations of 60 second blocks with EEGLAB `boundary` events marking the joins (Delorme and Makeig, 2004); we segmented each run into nonoverlapping 10 second windows and discarded any window intersecting a join to avoid edge transients. *Montage harmonization:* we used a canonical 59-channel 10–10 subset (Oostenveld and Praamstra, 2001); for each participant we applied the same presence mask (no interpolation) to the data columns and to the dictionary rows, preserving identical channel order across methods.

Participants with at least one valid EC window and one valid EO window after these steps were retained, yielding a final sample of *N* = 202. We also evaluated 32- and 19-channel subsets using the same masking strategy. Full channel lists and windowing details are provided in the Supplement (Section S1).

#### Time–frequency scalp topographies

Each EO/EC run was segmented into non-overlapping 10 s windows; for each window we computed Morlet-wavelet power on a logarithmic grid from 2–30 Hz (*F* =15 frequencies), which balances resolution at low vs. high frequencies (Cohen, 2019). For each frequency, sensor power was averaged over the 10 second window to yield one length-*M* topography. Stacking across windows and frequencies produced a subject-level matrix

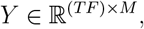

where *T* is the number of time windows, *F* the number of frequencies, and *M* the number of retained channels (after canonical masking). We also formed band-aggregated matrices *Y*_band_ ∈ ℝ^(*T* ×4)×*M*^ (delta 2–4 Hz, theta 4–7 Hz, alpha 8–12 Hz, beta 13–30 Hz). We *z*-scored each row of *Y* (or *Y*_band_) (per time-frequency slice) across channels to emphasize spatial configuration rather than absolute power level (i.e., removing an offset across channels). Full ingestion and TFR details are provided in the Supplement (Section S1).

#### Template LB basis and whole-cortex modes

On each hemisphere of *FreeSurfer* fsaverage (Dale et al., 1999; Fischl, 2012), we computed Laplace–Beltrami (LB) eigenmodes using the cotangent finite-element discretization of

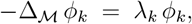

mass-normalized eigenvectors so that modes *ϕ*_*k*_ are orthonormal under the surface mass inner product (Meyer et al., 2003; Reuter et al., 2006); Δ_ℳ_ denotes the LB operator on the cortical surface ℳ. The modes *ϕ*_*k*_ are ordered from broad, brain-wide gradients (small *k*) to progressively finer structure (large *k*) (Pang et al., 2023). We retained *K*_hemi_=60 nontrivial modes per hemisphere.

The global sign of each eigenmode is arbitrary. To obtain bilaterally consistent whole-cortex modes, we aligned hemispheres by reflecting the right hemisphere (RH) across the mid-sagittal plane and mapping mirrored RH vertices to the left hemisphere (LH) via nearest-neighbor (NN) correspondence in 3D source space. Denoting this alignment by *T*_*R*→*L*_ = NN_*L*_◦Mirror (and writing 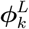 and 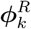 for the *k*-th LH/RH modes), we first define a global 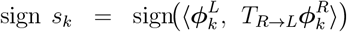 and then concatenate 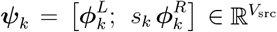, to obtain whole-cortex bilaterally consistent modes in source space. A step-by-step schematic is provided in Algorithm 1 (Supplement Section S2). This procedure yields *K* whole-cortex LB modes used in downstream sensor-space mapping.

### Forward mapping to sensor space (lead-field–mapped cortical harmonics)

Using MNE–Python, we defined a cortical *source space* on *fsaverage* with an icosahedral grid (ico4; ∼2,500 vertices per hemisphere, *V*_src_ ≈ 5,100 source vertices total) (Gramfort et al., 2014). The whole-cortex LB modes ***ψ***_*k*_ were reindexed to the source-space vertex order and stacked as columns of 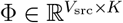. The EEG forward (lead-field) model was computed in MNE–Python using a three-layer BEM head model (scalp, skull, brain; conductivities 0.3/0.006/0.3 S/m) with fixed-orientation cortical dipoles aligned to the local surface normal (Vorwerk et al., 2014).Let 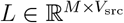 denote the resulting lead-field matrix. Mapping cortical LB modes to sensors gives a fixed, subject-agnostic sensor dictionary

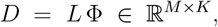

whose *k*th column is the scalp projection of the *k*th cortical harmonic.

Observed topographies were projected by ordinary least squares (OLS) onto the LB sensor subspace. Sensor locations followed the international 10–5 template (Oostenveld and Praamstra, 2001). These were reduced to our 59-channel subset, and for the 32- and 19-channel analyses we applied the same subject-specific channel mask to the rows of both *L* and *D*, and all methods operated on identical sensor subsets (see Supplement Section S1 for implementation details and defaults). Where source imaging seeks explicit cortical estimates, we instead assess representation quality directly in sensor space using a cortex-aligned dictionary, providing a complementary perspective to inverse solutions (Michel et al., 2004; Michel and Brunet, 2019).

#### Interpretability

Each column of *D* is the forward projection of a cortical harmonic with a known cortical spatial-frequency ordering and topology; thus low-order columns tend to emphasize broad, geometry-consistent patterns (e.g., posterior–anterior gradients) while remaining in sensor space (no subject-specific inverse solution required) (Pang et al., 2023). In practice, only the leading ∼ 10–20 harmonics contribute appreciably at the scalp, where higher-index cortical modes are strongly attenuated by the volume conductor and by montage sampling limits (after forward mapping and channel masking) (Nunez and Srinivasan, 2006; Hämäläinen et al., 1993; Vorwerk et al., 2014). An overview of the pipeline—from cortical LB modes on *fsaverage* to the sensor dictionary *D* = *L*Φ and OLS projection of each scalp topography *y* ∈ ℝ^*M*^ (per time-frequency slice)—is shown in Figure 1.

**Figure 1.**
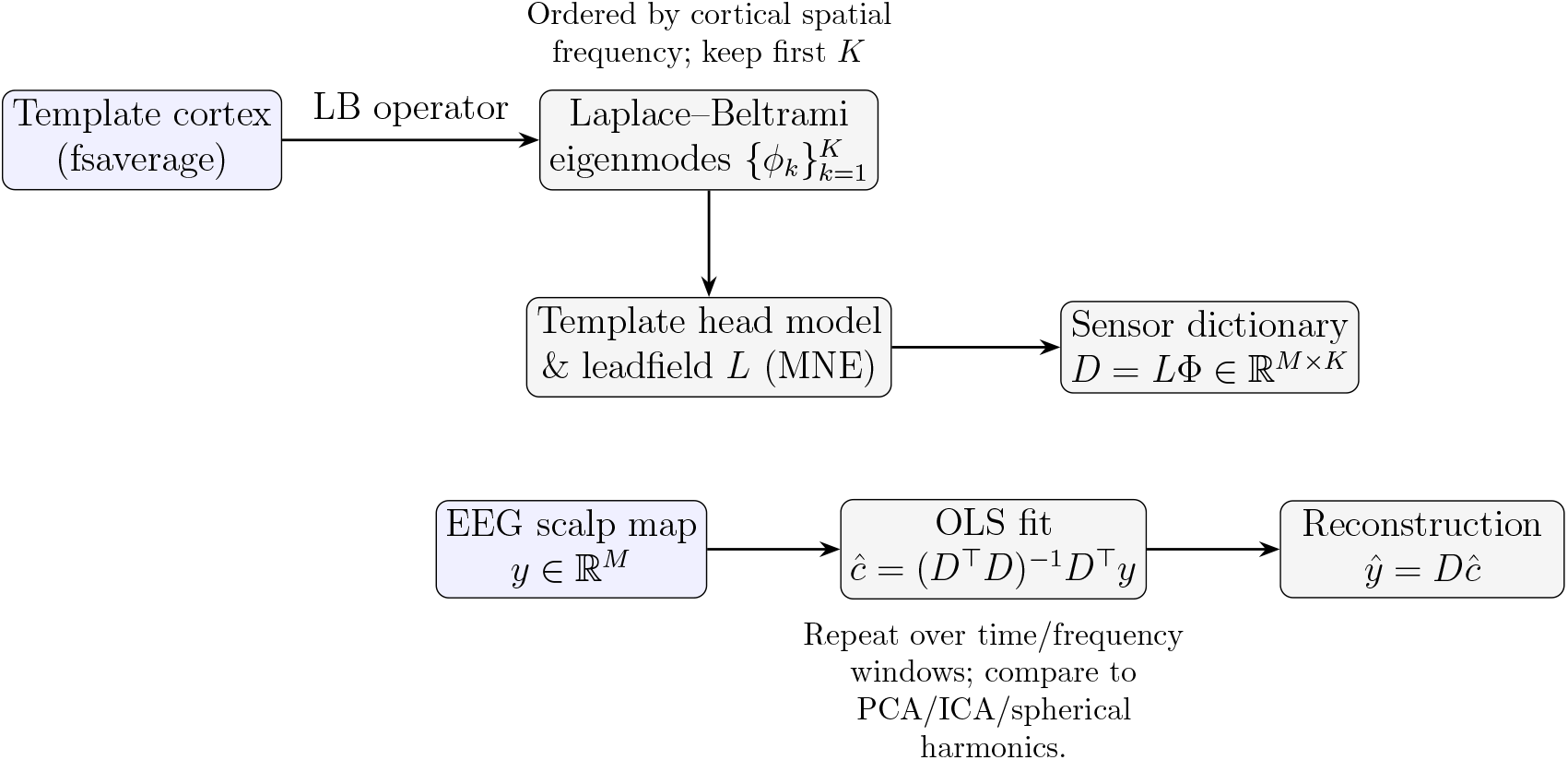
Pipeline for template LB representation of EEG. A standard cortical template provides Laplace–Beltrami (LB) eigenmodes; a template lead field maps these to a sensor-space dictionary *D*. Each EEG scalp map is expanded on *D* via OLS to obtain LB coefficients *ĉ* and a low-rank reconstruction *ŷ* ∈ ℝ^*M*^ for each time × frequency window (i.e., the same OLS operation is applied row-wise to *Y* ∈ ℝ^(*TF*)×*M*^).

### Representation and coefficient estimation (OLS span test)

We evaluated reconstruction efficiency of resting-state EEG (eyes–closed/open) for 59- , 32-, and 19–channel montages using OLS projections onto a basis that is orthonormalized in native order. Concretely, for each basis *B* (LB, SPH, PCA, ICA) we constructed an orthonormal column set *Q* ∈ ℝ^*M* ×*r*^ as follows. For LB and SPH, we applied thin QR without column pivoting to the first *K* columns of *B* (i.e., took the first *K* columns of the dictionary in their native order and re-orthonormalized this prefix via no-pivot QR), and dropped only numerically null directions using a relative tolerance *τ* = 10^−8^ on diag(*R*); this preserves the interpretable ordering (LB by cortical eigenvalue; SPH by spherical degree). For PCA, we used a thin SVD of the column-centered data matrix to obtain an orthonormal *Q*_PCA_ ordered by singular value (Jolliffe, 2002). For ICA, we fit FastICA on unit-variance–whitened rows to obtain the mixing-matrix topographies *A*, then orthonormalized *A* by QR (no pivot) to get *Q*_ICA_ in the algorithm’s component order (Hyvärinen and Oja, 2000; Delorme and Makeig, 2004). Projections used *Ŷ* = (*Y Q*)*Q*^T^, with *Y* ∈ ℝ^(*T F*)×*M*^ and each row *z*–scored across channels; truncation to *K* columns is done by replacing *Q* with *Q*_(:,1:*K*)_. For each subject and montage with *M* ∈ {59, 32, 19} channels, we swept *K* = 1, … , *r*, where *r* ≤ *M* is the effective rank after orthonormalization.

Efficiency was quantified by the variance explained *R*^2^(*K*) by *K* modes and the efficiency indices *K*_70_ and *K*_90_ (the smallest *K* with *R*^2^ ≥ 0.70 and 0.90); when a threshold was not attained by *K* = *r*, we reported the censored value at *K* = *r* together with an attainment indicator. For LB and SPH, per-mode reliability was assessed by ICC(3,1) of eyes–open/closed coefficients (band–mean OLS coefficients; *c* = *Y*_band_*Q*), using the same preprocessing and channel masks across the methods and montages (Shrout and Fleiss, 1979). To summarize, the comparator bases were:

- **PCA** (channel space): orthonormal principal axes *Q*_PCA_ from the subject’s *Y* rows after *column centering* (data-adaptive) (Jolliffe, 2002).
- **ICA** (FastICA): mixing matrix topographies *A* (unit-variance whiten), orthonormalized to *Q*_ICA_ (Hyvärinen and Oja, 2000; Delorme and Makeig, 2004).
- **SPH**: real spherical harmonics *B*_SPH_(*θ, ϕ*) evaluated at electrode locations, orthonormalized in degree-major order to *Q*_SPH_ (Perrin et al., 1989).

#### Initial dimensionality and ordering of modes

For LB we construct 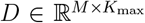 with *K*_max_ = 60 and then mask to the subject’s *M* retained channels, evaluating *K* ≤ *M* in native order. For SPH we generate at least *M* real spherical-harmonic columns (degree-major); for PCA the number of orthonormal channel PCs is at most min {*M*, rows(*Y*) }; and for ICA we cap the fitted components by min {*M*, rank(*Y*) }, restricting all methods to *K* ≤ *M* . We preserve the native, interpretable order within each basis: LB by increasing cortical eigenvalue (spatial frequency) of the source-space modes before forward mapping; SPH by increasing spherical degree (*ℓ* = 0, 1, 2, …) with meaningful checkpoints only at *K* = (*ℓ* + 1)^2^; PCA by descending singular values (variance captured); and ICA by the component order returned by whitening+FastICA (after orthonormalizing the mixing topographies).

For SPH, within a fixed spherical degree *ℓ*, the 2*ℓ* + 1 real spherical harmonics have no unique native ordering (arbitrary within-degree ordering). Therefore, for SPH we compute *R*^2^ and thresholds only at degree-block checkpoints *K* ∈ {(*ℓ* + 1)^2^} = {1, 4, 9, 16, 25, … .} For PCA/ICA/LB, we compute thresholds from the full *R*^2^(*K*) curves, whereas comparisons to SPH are made on these degree-block checkpoints.

### Evaluation metrics

#### (1) Variance explained (efficiency)

For each basis size *K* and method, we compute explained variance *row-wise* for each time–frequency topography *y* ∈ ℝ^*M*^ (one row of *Y* ∈ ℝ^(*T F*)×*M*^):

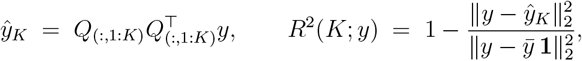

where 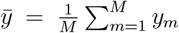 . We average *R*^2^(*K*; *y*) over the time–frequency slices within subject (*s*) and condition (*c*) to obtain 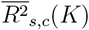; group summaries report 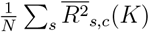 along with 95% bootstrap confidence intervals (CIs), resampling subjects. Efficiency is summarized by the group mean *R*^2^(*K*) curve and by the efficiency indices *K*_70_ and *K*_90_. We also report normalized indices (*K*_70_*/M, K*_90_*/M*) for cross-montage comparability and apply right-censoring at *K* = *r* with attainment indicators.

#### (2) Across-condition reliability (EO vs. EC)

We assess stability by ICC(3,1) (Shrout and Fleiss, 1979). For LB and SPH (bases with fixed modes), we compute OLS coefficients *c* = *Q*^T^*Y* and aggregate within canonical bands (*δ, θ, α, β*) to obtain band-mean coefficients; we then compute ICC(3,1) per mode and band comparing EO vs. EC within subject, with bootstrap confidence intervals (see Supplement Section S3 for the ICC formula).

#### (3) Montage robustness

Channel order is standardized to a 59-channel subset derived from `standard_1005` (excluding non-EEG sensors; the list of the 59-channel subset in Supplement Section S1) (Oostenveld and Praamstra, 2001). Robustness to spatial sampling is evaluated by repeating (1)–(2) on 32- and 19-channel subsets. We report *R*^2^(*K*) over *K* and distributions of *K*_70_ and *K*_90_ (*K*_70_*/M* and *K*_90_*/M*) across montages.

#### (4) Computational efficiency

We quantified runtime cost as wall-clock seconds per 1,000 topographic rows for the OLS projection path, timing each method under the 59-channel montage and a fixed target dimension (*K*=20). For every subject and condition, the pipeline consisted of (i) constructing an orthonormal basis *Q* in the method’s native order (LB and SPH: thin QR without column pivoting (prefix per *K*) on the masked dictionary/design; PCA: SVD of column-centered *Y*; ICA: SVD-orthonormalization of the mixing-matrix topographies), and (ii) projecting *Y* via *Ŷ* = (*Y Q*)*Q*^T^. Times were averaged across five repeats and summarized as the median seconds per 1,000 rows across subjects (*N* = 202). After basis formation, the per-row projection cost for LB scales as 𝒪 (*MK*) and is typically far lower than ICA fitting.

### Statistical analysis

Group summaries use percentile bootstrap confidence intervals (resampling subjects, *B* = 2000). For *R*^2^(*K*) we bootstrap the group-mean curve over a grid of *K*; for thresholds *K*_70_ and *K*_90_ we report attainment rates and the mean (and normalized) *K*_70_ and *K*_90_ among attainers (with CIs). For reliability we report ICC(3,1) per mode×band with bootstrap CIs. The subject-level resampling procedure is detailed in the Supplement (Section S4).

### Reproducibility

We provide reproducible Python code to: (i) build the sensor dictionary *D* = *L*Φ_sym_ on *fsaverage* (ico4) using MNE–Python/BEM; (ii) construct SPH bases in sensor geometry; run OLS span tests for LB/PCA/ICA/SPH with channel alignment and montage subsets; and (iv) generate outputs for group summaries (bootstrap mean *R*^2^(*K*) with CIs), *K*_90_ attainment/values, and mode-wise ICC tables. Canonical channel lists (59/32/19) are included. Implementation details are provided in the Supplement (Section S5).

## Results

We compared four sensor–space bases—lead field–mapped cortical harmonics (LB), spherical harmonics (SPH), PCA, and ICA—using OLS projections on resting EEG (eyes closed, EC; eyes open, EO) across 59-, 32-, and 19-channel montages.

### Representational efficiency (59 channels)

Figure 2 illustrates *R*^2^(*K*) for the early-*K* regime. At *K*=4 (EC), LB already explains 0.565 [95% CI: 0.543, 0.585] of variance, clearly ahead of SPH 0.437 [0.417, 0.458], while PCA/ICA capture little (PCA 0.075 [0.067, 0.085]; ICA 0.078 [0.070, 0.085]). The same ordering holds for EO at *K*=4 (LB 0.441 [0.419, 0.464]; SPH 0.356 [0.332, 0.380]; PCA 0.104 [0.096, 0.112]; ICA 0.098 [0.090, 0.107]). By *K*=9 the two fixed bases are close (EC: LB 0.720 vs SPH 0.716); by *K*=16 both are high (EC: LB 0.838, SPH 0.851); and by *K*=25 they effectively converge (EC: LB 0.916, SPH 0.915), while PCA approaches unity as *K* nears the montage rank (Table 1). Overall, LB delivers the steepest early rise and the best fidelity at small *K* (up to ∼ 9). Extended LB checkpoints for *R*^2^(*K*) at *K* = 5, 10, 15, 20, 25, … for both EC and EO are provided in Table 2.

**Table 1:**
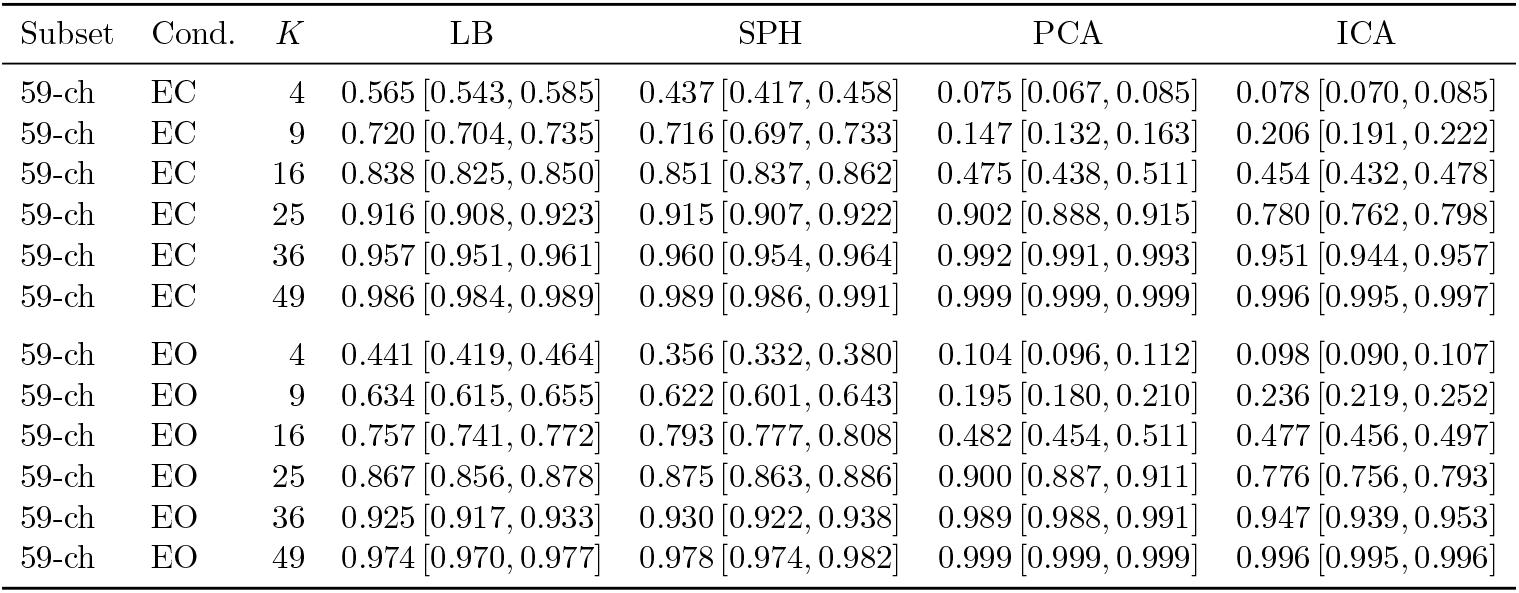
Group-mean explained variance *R*^2^(*K*) (bootstrap 95% CIs over subjects; *N* =202) for the 59-channel montage at selected basis sizes *K* ∈ {4, 9, 16, 25, 36, 49 }, shown separately for EC and EO. LB yields higher early-*K* fidelity (larger *R*^2^) than SPH/PCA/ICA (e.g., *K*=4).

**Table 2:**
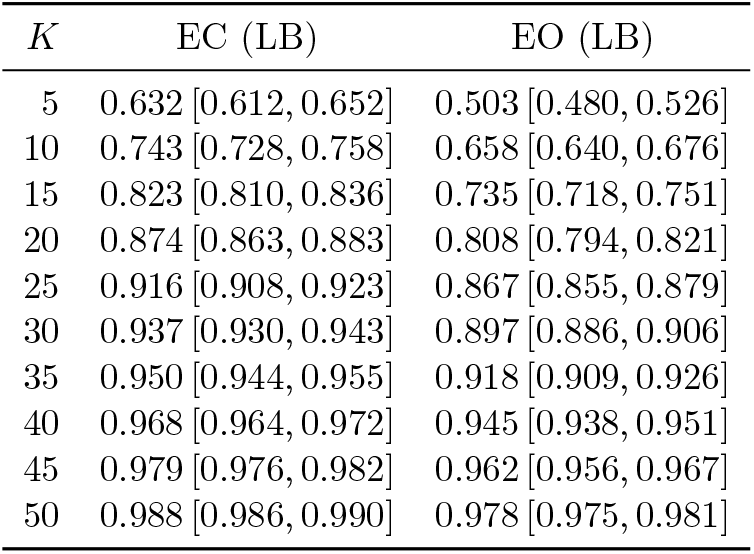
Extended LB checkpoints (59-channel montage): group-mean explained variance *R*^2^(*K*) with 95% bootstrap CIs for EC and EO over the leading *K* modes, where *K*=5, 10, … , 50.

| $K$ | EC (LB) | EO (LB) |
| --- | --- | --- |
| 5 | 0.632 [0.612, 0.652] | 0.503 [0.480, 0.526] |
| 10 | 0.743 [0.728, 0.758] | 0.658 [0.640, 0.676] |
| 15 | 0.823 [0.810, 0.836] | 0.735 [0.718, 0.751] |
| 20 | 0.874 [0.863, 0.883] | 0.808 [0.794, 0.821] |
| 25 | 0.916 [0.908, 0.923] | 0.867 [0.855, 0.879] |
| 30 | 0.937 [0.930, 0.943] | 0.897 [0.886, 0.906] |
| 35 | 0.950 [0.944, 0.955] | 0.918 [0.909, 0.926] |
| 40 | 0.968 [0.964, 0.972] | 0.945 [0.938, 0.951] |
| 45 | 0.979 [0.976, 0.982] | 0.962 [0.956, 0.967] |
| 50 | 0.988 [0.986, 0.990] | 0.978 [0.975, 0.981] |

**Figure 2.**
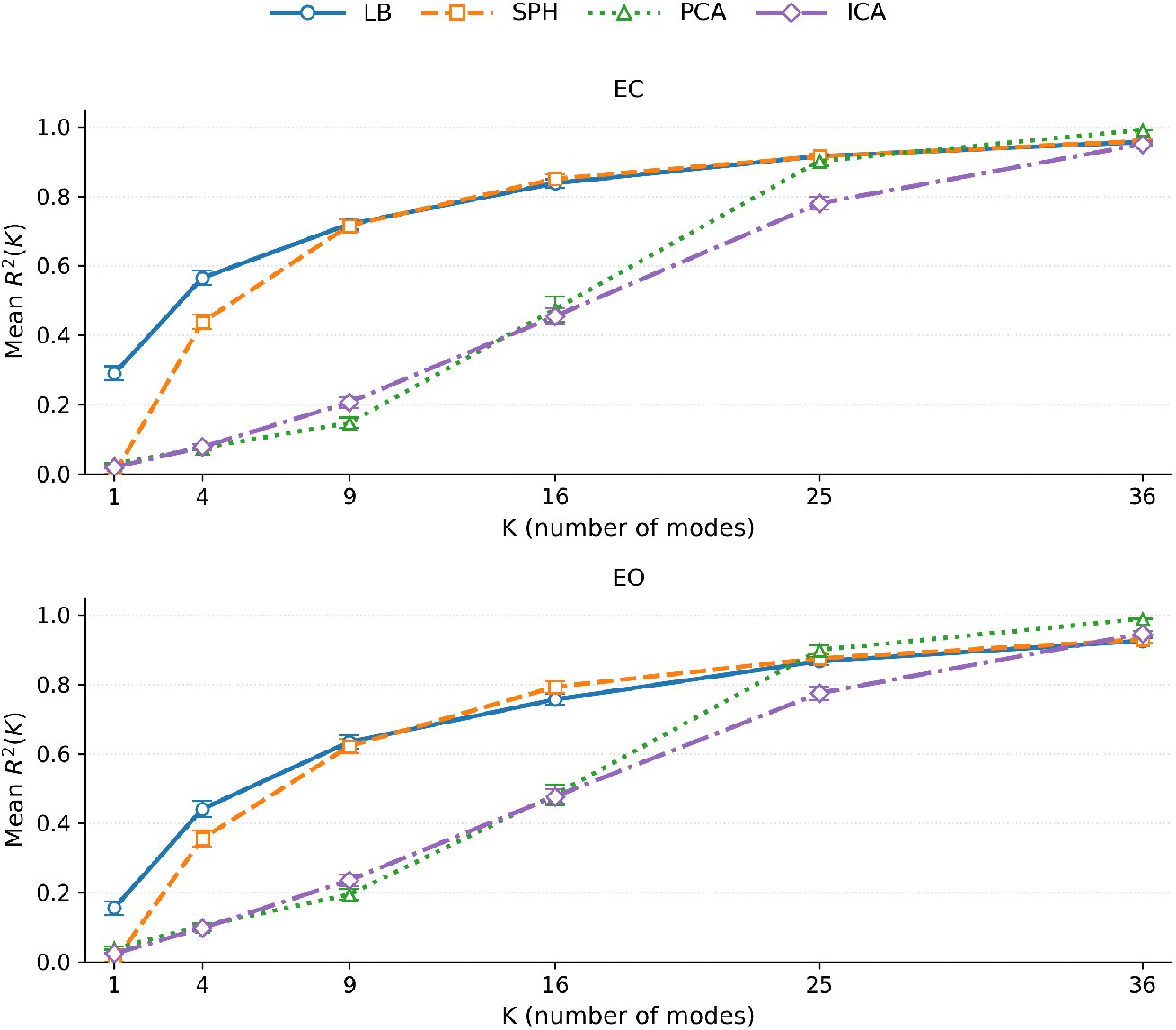
Efficiency *R*^2^(*K*) at 59 channels. Group-mean explained variance *R*^2^(*K*) at *K* ∈ {1, 4, 9, 16, 25, 36 }, shown separately for eyes-closed (EC; top panel) and eyes-open (EO; bottom panel), with bootstrap 95% CIs over subjects (*N* =202). LB rises steeply and leads at very small *K* (1–4), SPH catches up by *K* ≈ 9 and then tracks LB closely, while PCA/ICA lag at small *K* but approach unity near *K* = 25–36. The same trend holds for both EC and EO.

### Montage robustness (59 → 32 → 19)

The early-*K* advantage of LB persists under channel reduction to 32 and 19 (see Table 3). At *K*=4, LB exceeds SPH in both EC and EO for the 32- and 19-channel subsets (e.g., 32-ch EC: LB 0.606 [0.584, 0.628] vs SPH 0.478 [0.456, 0.501]; 19-ch EC: LB 0.587 [0.563, 0.610] vs SPH 0.491 [0.468, 0.515]), indicating that a few cortex-aligned modes still capture the dominant scalp structure even with sparse sampling. By *K*=9, the fixed bases are essentially tied in 32-ch (EC: LB 0.770 vs. SPH 0.768; EO: LB 0.682 vs. SPH 0.668) and closely matched in 19-ch (EC: LB 0.797 vs. SPH 0.804; EO: LB 0.723 vs. SPH 0.715). At *K*=16, both fixed bases achieve high *R*^2^ across montages, while PCA continues to climb toward unity as *K* approaches the montage rank. Because LB is ordered by cortical spatial frequency and the forward model already attenuates fine detail, and because down-sampling primarily removes that fine detail, the first few LB modes remain well matched to the large-scale patterns resolvable at the scalp.

**Table 3:** Montage robustness of *R*^2^(*K*) for 32- and 19-channel subsets (EC/EO). Entries are group-mean *R*^2^(*K*) with 95% bootstrap CIs. Reported *K* values are {4, 9, 16, 25} for 32-ch and {4, 9, 16} for 19-ch (limited by *M*). LB retains its early-*K* advantage under channel down-sampling.

| Subset | Cond. | $K$ | LB | SPH | PCA | ICA |
| --- | --- | --- | --- | --- | --- | --- |
| 32-ch | EC | 4 | 0.606 [0.584, 0.628] | 0.478 [0.456, 0.501] | 0.081 [0.071, 0.092] | 0.086 [0.078, 0.095] |
| 32-ch | EC | 9 | 0.770 [0.754, 0.785] | 0.768 [0.749, 0.786] | 0.171 [0.154, 0.191] | 0.254 [0.233, 0.274] |
| 32-ch | EC | 16 | 0.883 [0.873, 0.893] | 0.896 [0.884, 0.906] | 0.629 [0.593, 0.666] | 0.575 [0.549, 0.601] |
| 32-ch | EC | 25 | 0.969 [0.965, 0.973] | 0.966 [0.960, 0.970] | 0.987 [0.985, 0.989] | 0.929 [0.916, 0.940] |
| 32-ch | EO | 4 | 0.469 [0.444, 0.494] | 0.373 [0.348, 0.399] | 0.113 [0.104, 0.123] | 0.118 [0.108, 0.129] |
| 32-ch | EO | 9 | 0.682 [0.663, 0.700] | 0.668 [0.647, 0.688] | 0.222 [0.205, 0.239] | 0.288 [0.269, 0.308] |
| 32-ch | EO | 16 | 0.819 [0.806, 0.831] | 0.854 [0.841, 0.866] | 0.643 [0.613, 0.670] | 0.602 [0.578, 0.625] |
| 32-ch | EO | 25 | 0.955 [0.950, 0.960] | 0.954 [0.949, 0.959] | 0.985 [0.982, 0.988] | 0.928 [0.919, 0.936] |
| 19-ch | EC | 4 | 0.587 [0.563, 0.610] | 0.491 [0.468, 0.515] | 0.087 [0.078, 0.099] | 0.098 [0.088, 0.109] |
| 19-ch | EC | 9 | 0.797 [0.783, 0.810] | 0.804 [0.785, 0.821] | 0.195 [0.174, 0.217] | 0.289 [0.266, 0.311] |
| 19-ch | EC | 16 | 0.968 [0.963, 0.973] | 0.960 [0.953, 0.965] | 0.910 [0.884, 0.935] | 0.869 [0.846, 0.890] |
| 19-ch | EO | 4 | 0.447 [0.423, 0.471] | 0.391 [0.365, 0.416] | 0.123 [0.115, 0.133] | 0.122 [0.112, 0.133] |
| 19-ch | EO | 9 | 0.723 [0.706, 0.741] | 0.715 [0.696, 0.736] | 0.250 [0.231, 0.269] | 0.349 [0.326, 0.371] |
| 19-ch | EO | 16 | 0.941 [0.932, 0.950] | 0.938 [0.928, 0.947] | 0.923 [0.903, 0.941] | 0.881 [0.862, 0.900] |

### Attainment and efficiency indices (K_70_, K_90_)

Figures 3–4 show subject-level boxplots for the number of modes needed to reach *R*^2^ ≥ 0.70 (Figure 3) and *R*^2^ ≥ 0.90 (Figure 4), respectively, at 59 channels (attainment counts printed above each box); Table 4 reports corresponding means for *K*_70_ and *K*_90_ among attainers and their normalized values (*K/M*) across all montages and conditions, along with 95% bootstrap CIs. For *K*_70_ (see Figure 3), LB consistently requires the fewest modes and shows the tightest spread in both EC and EO, indicating a lower-dimensional representation to reach 70% variance. For *K*_90_ (Figure 4), LB remains efficient at 59-ch EC, requiring fewer modes than SPH and ICA and comparable to PCA (LB 22.6 (0.389*M*); SPH 25.3 (0.436*M*); PCA 23.2 (0.399*M*); ICA 30.2 (0.519*M*)). EO is uniformly harder (higher *K*_90_) across methods; PCA is lowest on average (PCA 24.2 (0.416*M*) vs. LB 29.3 (0.504*M*), SPH 31.4 (0.542*M*), ICA 30.4 (0.524*M*)), while LB preserves the early-*K* advantage. Normalized indices (*K/M*) show the same pattern for the 32- and 19-channel montages (Table 4).

**Table 4:** Efficiency indices among attainers across montages: mean *K*_70_ and *K*_90_ (and normalized *K/M*) with percentile bootstrap 95% CIs by montage, condition, and method (*N* =202) among “attainers.” An “attainer” is a subject for whom the threshold *R*^2^ ≥ 0.70 or 0.90 is reached by *K* ≤ *r* (effective rank). Right-censored cases are excluded from the means.

| Subset | Cond. | Method | $K_{70}$ | $K_{70}/M$ | $K_{90}$ | $K_{90}/M$ |
| --- | --- | --- | --- | --- | --- | --- |
| 59-ch | EC | LB | 8.604 [7.916, 9.332] | 0.149 [0.136, 0.161] | 22.569 [21.485, 23.560] | 0.389 [0.371, 0.408] |
| 59-ch | EC | SPH | 12.361 [11.698, 13.059] | 0.213 [0.201, 0.225] | 25.317 [23.970, 26.639] | 0.436 [0.413, 0.459] |
| 59-ch | EC | PCA | 19.347 [18.807, 19.936] | 0.333 [0.324, 0.343] | 23.168 [22.594, 23.763] | 0.399 [0.389, 0.410] |
| 59-ch | EC | ICA | 22.881 [22.322, 23.510] | 0.394 [0.384, 0.404] | 30.153 [29.440, 30.837] | 0.519 [0.508, 0.532] |
| 59-ch | EO | LB | 12.901 [12.000, 13.862] | 0.222 [0.207, 0.239] | 29.277 [28.005, 30.574] | 0.504 [0.483, 0.526] |
| 59-ch | EO | SPH | 15.644 [14.624, 16.713] | 0.270 [0.253, 0.288] | 31.448 [29.991, 32.876] | 0.542 [0.517, 0.566] |
| 59-ch | EO | PCA | 19.460 [18.995, 19.931] | 0.335 [0.327, 0.343] | 24.153 [23.673, 24.659] | 0.416 [0.408, 0.425] |
| 59-ch | EO | ICA | 22.446 [21.817, 23.079] | 0.387 [0.376, 0.398] | 30.406 [29.748, 31.084] | 0.524 [0.513, 0.536] |
| 32-ch | EC | LB | 6.807 [6.292, 7.332] | 0.217 [0.201, 0.234] | 16.797 [16.123, 17.431] | 0.535 [0.515, 0.556] |
| 32-ch | EC | SPH | 10.931 [10.371, 11.525] | 0.348 [0.330, 0.368] | 18.036 [17.263, 18.835] | 0.575 [0.550, 0.602] |
| 32-ch | EC | PCA | 16.371 [15.931, 16.797] | 0.522 [0.508, 0.535] | 19.109 [18.678, 19.515] | 0.609 [0.596, 0.622] |
| 32-ch | EC | ICA | 18.708 [18.228, 19.208] | 0.596 [0.579, 0.611] | 22.931 [22.485, 23.356] | 0.731 [0.716, 0.745] |
| 32-ch | EO | LB | 10.089 [9.421, 10.743] | 0.321 [0.300, 0.343] | 19.787 [19.178, 20.347] | 0.630 [0.611, 0.648] |
| 32-ch | EO | SPH | 13.396 [12.703, 14.134] | 0.427 [0.405, 0.450] | 21.212 [20.543, 21.891] | 0.677 [0.654, 0.700] |
| 32-ch | EO | PCA | 16.723 [16.376, 17.089] | 0.533 [0.521, 0.544] | 19.624 [19.282, 20.000] | 0.625 [0.614, 0.636] |
| 32-ch | EO | ICA | 18.193 [17.747, 18.654] | 0.580 [0.566, 0.595] | 23.292 [22.886, 23.713] | 0.742 [0.728, 0.754] |
| 19-ch | EC | LB | 6.371 [6.015, 6.752] | 0.344 [0.323, 0.365] | 12.550 [12.198, 12.891] | 0.676 [0.656, 0.694] |
| 19-ch | EC | SPH | 9.876 [9.418, 10.313] | 0.532 [0.508, 0.558] | 13.808 [13.346, 14.269] | 0.745 [0.719, 0.769] |
| 19-ch | EC | PCA | 13.955 [13.653, 14.248] | 0.752 [0.736, 0.768] | 14.812 [14.564, 15.074] | 0.798 [0.786, 0.811] |
| 19-ch | EC | ICA | 14.228 [13.931, 14.530] | 0.767 [0.751, 0.783] | 15.871 [15.658, 16.084] | 0.855 [0.844, 0.866] |
| 19-ch | EO | LB | 8.723 [8.272, 9.178] | 0.469 [0.445, 0.492] | 14.089 [13.738, 14.431] | 0.759 [0.741, 0.777] |
| 19-ch | EO | SPH | 11.530 [11.035, 12.086] | 0.621 [0.593, 0.648] | 15.279 [14.939, 15.576] | 0.826 [0.807, 0.842] |
| 19-ch | EO | PCA | 14.074 [13.812, 14.327] | 0.758 [0.744, 0.771] | 15.262 [15.050, 15.465] | 0.822 [0.811, 0.832] |
| 19-ch | EO | ICA | 13.653 [13.366, 13.965] | 0.735 [0.718, 0.752] | 15.738 [15.515, 15.965] | 0.847 [0.835, 0.858] |

**Figure 3.**
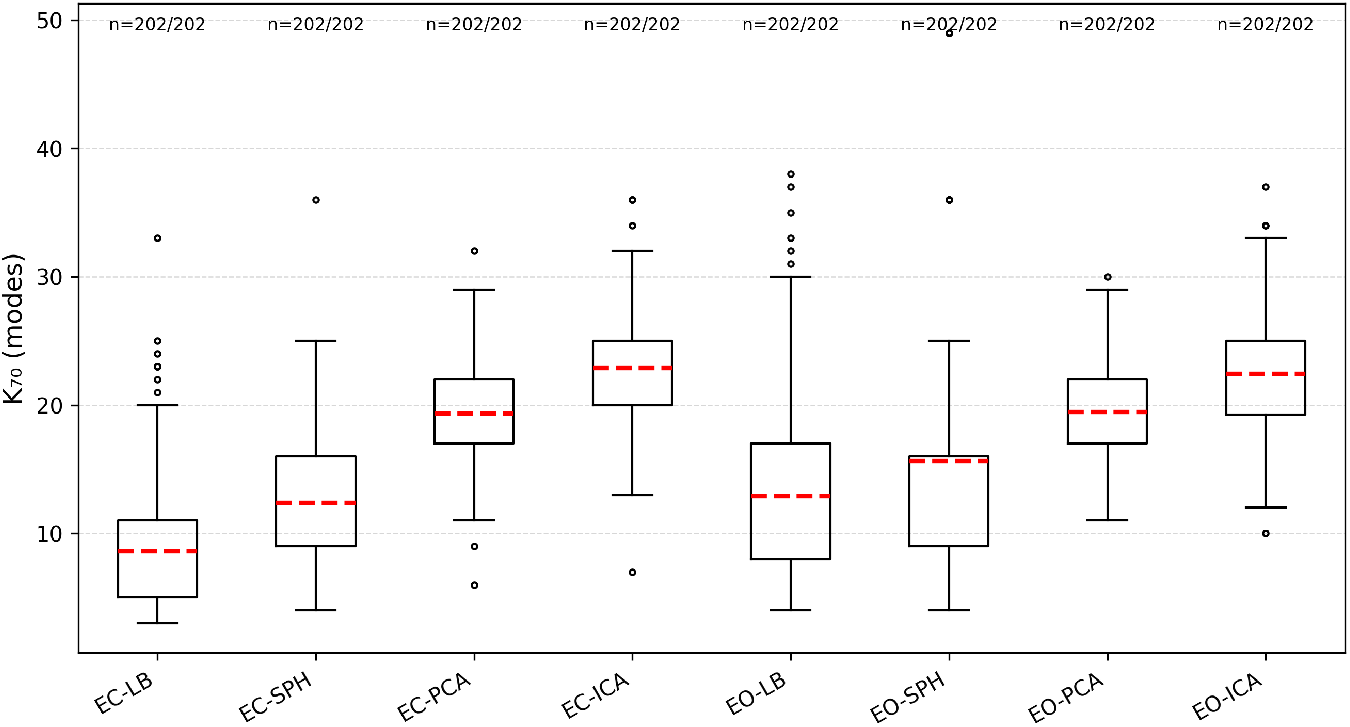
Threshold efficiency at 59 channels: *K*_70_ (number of modes needed to reach *R*^2^ ≥ 0.70). Boxplots show the per-subject number of modes required to reach *R*^2^ ≥ 0.70 (*K*_70_), for each method (LB/SPH/PCA/ICA) and condition (EC/EO). Red dashed lines mark the group mean; boxes show mean and IQR; whiskers extend to 1.5 IQR. Labels above boxes give the attainment count *n/N* (all 202*/*202). LB has the smallest central tendency for *K*_70_ in both EC and EO, indicating fewer modes are needed to reach 70% variance than SPH, PCA, or ICA.

**Figure 4.**
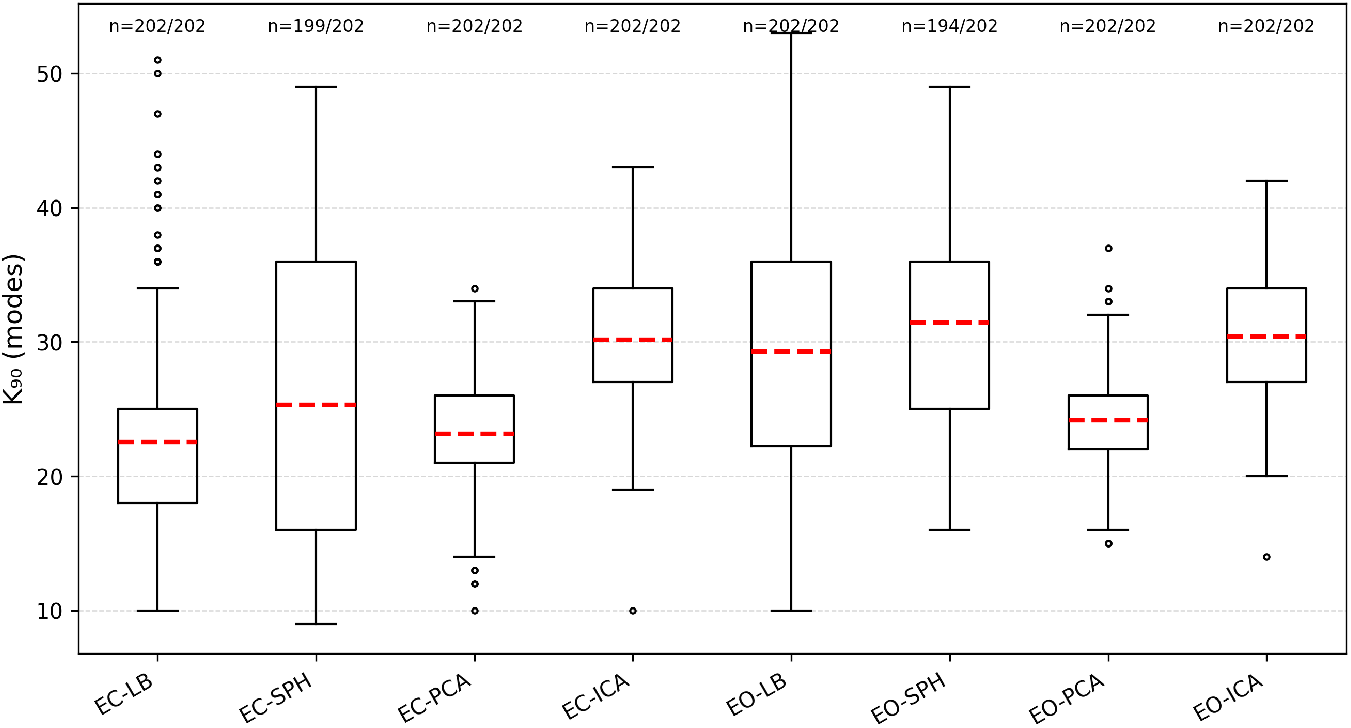
Threshold efficiency at 59 channels: *K*_90_ (number of modes needed to reach *R*^2^ ≥ 0.90). Boxplots show the per-subject number of modes required to reach *R*^2^ ≥ 0.90 (*K*_90_), for each method (LB/SPH/PCA/ICA) and condition (EC/EO). Red dashed lines mark the group mean; boxes show the median and IQR; whiskers extend to 1.5 IQR.

### Reliability (EO–EC ICC)

Per-mode EO–EC reliability was similar in magnitude for LB and SPH across all bands (Table 5). Both bases yield mid-0.4 mean ICCs with overlapping CIs; small band-specific differences appear (LB is slightly higher in *α/θ*, SPH slightly higher in *β*) but the overall pattern is comparable. In terms of coverage, nearly all modes exceed ICC ≥ 0.3, and roughly a third exceed ICC ≥ 0.5 for both bases, indicating that a substantial portion of the low–mid modes carry stable coefficients across EO and EC. For SPH, we also report degree-aggregated ICCs that combine orders within each spherical degree (Table S3). This aggregation is helpful for viewing reliability by spatial scale, though it compresses within-degree variability; all ICCs *<* 0.5 for this SPH degreeaggregation. Mode-wise tables (Supp. Tables S1, S2) show broadly similar reliability profiles for LB and SPH across many low–mid modes.

**Table 5:** Across-condition reliability coverage at *K* = 20 (59-ch): fraction of modes exceeding ICC thresholds (0.3, 0.5) and mean ICC(3,1) with 95% bootstrap CIs, computed from EO vs. EC band-mean OLS coefficients. LB and SPH show comparable mode-wise reliability across *α, β, δ*, and *θ* bands.

| Method | Band | % ICC $\geq 0.3$ | % ICC $\geq 0.5$ | Mean ICC [95% CI] |
| --- | --- | --- | --- | --- |
| LB | $\alpha$ | 100 | 40 | 0.472 [0.433, 0.510] |
| SPH | $\alpha$ | 95 | 30 | 0.470 [0.436, 0.500] |
| LB | $\beta$ | 100 | 35 | 0.472 [0.437, 0.507] |
| SPH | $\beta$ | 95 | 40 | 0.474 [0.438, 0.506] |
| LB | $\delta$ | 95 | 40 | 0.466 [0.424, 0.507] |
| SPH | $\delta$ | 95 | 30 | 0.462 [0.429, 0.492] |
| LB | $\theta$ | 100 | 40 | 0.478 [0.437, 0.518] |
| SPH | $\theta$ | 95 | 30 | 0.469 [0.434, 0.500] |

### Spatial exemplars with high reliability

Figure 5 illustrates LB mode 1 and three low–mid modes (4, 5, 8) selected by *α*-band EO–EC reliability (ICC(3,1) *>* 0.5) among the leading *K* = 10 modes. Point estimates (95% bootstrap CIs) are: Mode 4 0.560 [0.436, 0.663], Mode 5 0.553 [0.401, 0.671], Mode 8 0.607 [0.508, 0.696]; Mode 1 is shown for reference (0.472 [0.361, 0.577]). In Figure 5, cortical panels (left) display the template LB patterns on *fsaverage* (LH shown); sensor panels (right) show the corresponding forward-projected topographies. These organization patterns (e.g., posterior–anterior gradients in Mode 1, posterior *α* dominance) are consistent with classic findings on occipital alpha topography and its cortical generators (Nunez et al., 2001; Barzegaran et al., 2017; Michel and Murray, 2012). Because LB mode identities are shared across participants, the LB basis provides a reproducible coordinate system for reporting effect topographies (e.g., “LB mode 8 in *α*”). As the mode index increases, cortical spatial frequency increases (Mode 4 is coarser than 5 and 8); this coarse-to-fine ordering is useful and compatible with the idea that large-scale gradients dominate scalp-resolvable variance, with finer features attenuated by the volume conductor.

**Figure 5.**
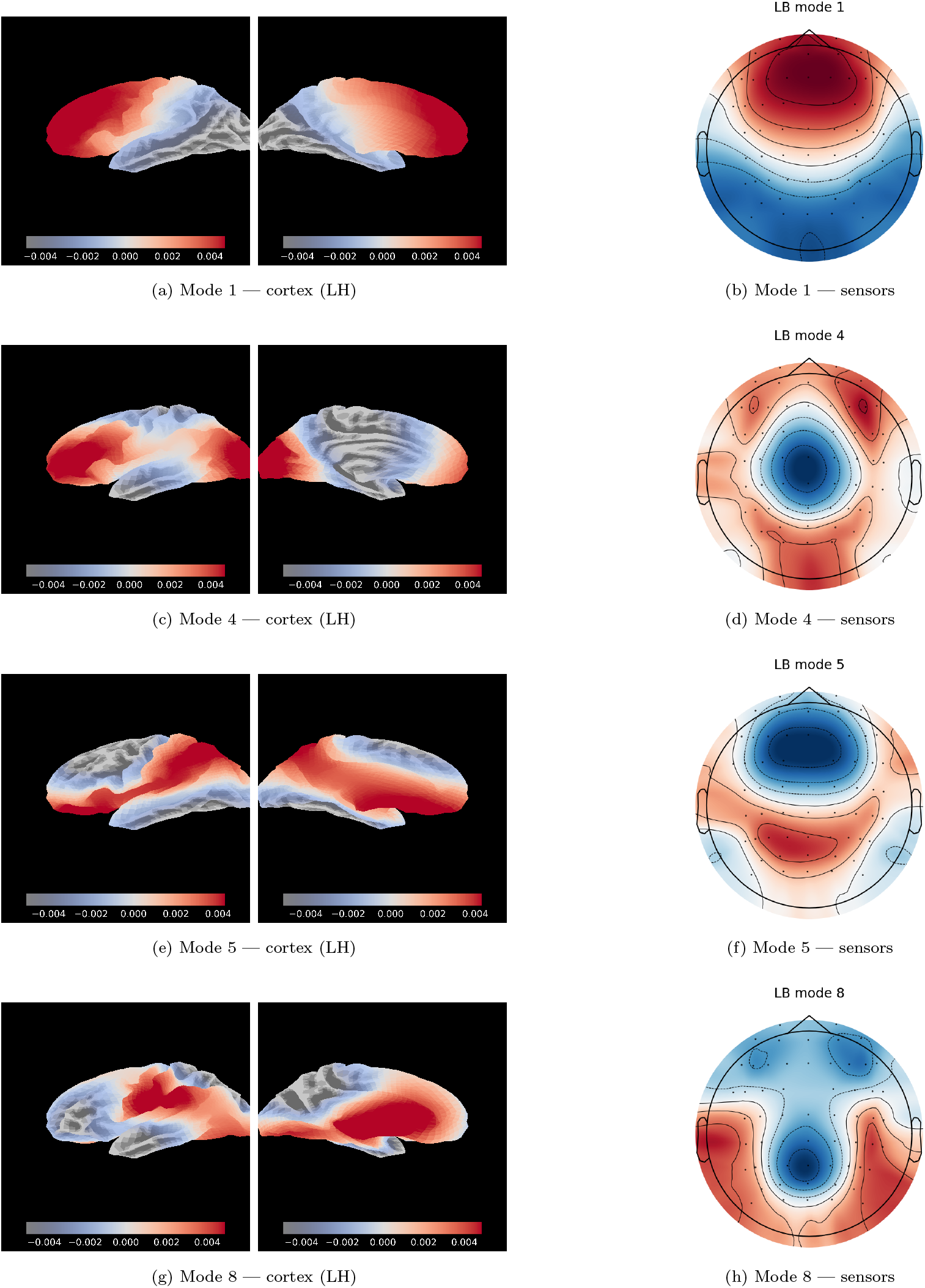
LB spatial exemplars. Modes were selected by *α*-band EC–EO ICC(3,1) *>* 0.5 (Mode 1 included as the leading mode). Mean ICC(3,1) [95% bootstrap CI] for the *α* band: Mode 1 0.472 [0.361, 0.577], Mode 4 0.560 [0.436, 0.663], Mode 5 0.553 [0.401, 0.671], Mode 8 0.607 [0.508, 0.696]. Left: cortical maps on *fsaverage* (left hemisphere shown; whole-cortex modes are constructed to be bilaterally consistent). Right: corresponding sensor-space projections. *Note*. Mild scalp asymmetries arise from the head model, montage sampling, and slight inter-hemispheric anatomical differences.

### Computational efficiency

Supplementary Table S4 reports projection throughput on the 59-channel montage at *K*=20, including per-subject basis orthonormalization and the OLS map *Y* → (*Y Q*)*Q*^T^. Median time per 1,000 band-averaged rows was: LB 1.48–1.50 ms, SPH 3.40–3.43 ms, PCA 4.90–4.93 ms, and ICA 100.6–108.9 ms (EC–EO). Thus, LB is the fastest among the tested bases: about 2.3× faster than SPH, 3.3× faster than PCA, and ∼ 70× faster than ICA, with essentially identical timings in EC and EO. These results indicate that geometry-aligned LB projection attains near-trivial computational cost while preserving the early-*K* fidelity advantages documented above.

## Discussion

### Summary of findings

We asked whether scalp EEG can be represented in a compact, geometry-aligned subspace spanned by cortical Laplace–Beltrami (LB) modes mapped to sensors. Across eyes-closed/open (EC/EO) resting EEG and three montages (59/32/19 channels), LB captured more variance with fewer modes—higher early-*K R*^2^ and smaller *K*_70_, *K*_90_ (and *K/M*)—than SPH. These trends held under channel down-sampling, indicating robustness to reduced spatial sampling. LB coefficients also showed mode-wise EO–EC reliability comparable to SPH. Together, the results support a cortex-aligned, low-dimensional account of scalp EEG that requires neither subject-specific MRI nor full source inversion.

### Why geometry helps

Unlike sensor-geometry bases, LB modes originate on the cortical surface and are ordered by intrinsic spatial frequency. Forward projection through a realistic head model preserves both 1) the multiscale ordering and 2) anatomical provenance in the sensor dictionary *D*, enabling statements such as “mid-order modes carry *β* over sensorimotor regions” in a physically meaningful coordinate system. As the skull and scalp act as a spatial low-pass filter, scalp EEG predominantly reflects these large-scale cortical patterns. In practice, this geometry alignment appears to (i) concentrate variance into the first ∼ 10–20 modes (steeper *R*^2^(*K*)), (ii) yield many modes with non-trivial ICC coverage at distinctive spatial scales, and (iii) avoid over-interpretation of fine detail that scalp EEG is unlikely to resolve.

### Relation to conventional decompositions

PCA and ICA are powerful data-adaptive tools, but they lack an anatomical frame: their axes are sample-specific and require post-hoc alignment for cross-subject comparability. In span tests, PCA predictably approaches unity as *K* nears the montage rank, yet in the compression regime that matters (small-moderate *K*) it typically underperforms LB, reflecting a less favorable variance-complexity trade-off. ICA excels at source separation and artifact removal, but as a representational basis it performs poorly here, with high *K*_70_ and low *R*^2^(*K*) at small-moderate *K*. SPH offers a fixed sensor-space basis, but orders within each spherical degree are rotationally ambiguous and carry no cortical correspondence, limiting interpretability. Our results align with the view that analyzing the scalp electric field in anatomically grounded coordinates improves interpretability over channel-wise or purely data-adaptive bases (Michel and Murray, 2012). All comparisons used identical preprocessing, channel masks, and *K* ≤ *M* (with SPH evaluated at degree-block checkpoints to respect its native structure).

### Interpretability and neurophysiology

Low-order LB modes visualized on cortex and at the scalp show canonical anterior-posterior and sensorimotor organization consistent with prior EEG/MEG literature (Pfurtscheller and Lopes da Silva, 1999; Hari and Puce, 2017). Because LB coefficients are anatomy-anchored and share mode identities across participants, they support reproducible reporting and facilitate direct links to MRI-based analyses, bridging sensor-space EEG with cortical geometry rather than treating them as separate domains.

### Practical implications

Once the template dictionary *D* = *L*Φ is precomputed, LB coefficients can be obtained at a near-trivial computational cost, scaling as 𝒪 (*MK*), without unstable source inversion. Because the basis is fixed and shared across subjects, it naturally supports cross-subject summaries, bootstrap uncertainty quantification, and downstream modeling (e.g., regression/decoding) with interpretable, geometry-aware covariates.

### Limitations

First, we use a template (fsaverage) rather than individual cortical geometry. While this choice enables subject-agnostic coefficients, it neglects individual anatomical variation; future work can incorporate subject-specific surfaces or morph the template modes to subject space. Second, forward models (conductivities, skull thickness) are approximate; although our results were robust across montages, improved head modeling could refine sensor-space projections, Third, we focused on resting topographies and band-mean coefficients; extensions to task-evoked patterns and time-resolved coefficients will further validate generality. Finally, we emphasized LB versus SPH/PCA/ICA span properties; integrating other cortical bases (e.g., network- or connectivity-aligned) is an interesting direction.

## Conclusion

A compact set of forward-projected cortical eigenmodes offers an efficient, anatomically interpretable basis for sensor-space resting-state EEG. In the operating regime that matters for compression and interpretability (small-moderate *K*), the LB basis attains higher fidelity with fewer modes and robust EO/EC reliability, while remaining simple to compute and reproduce. This geometry-aligned representation should facilitate scalable, multimodal, and longitudinal EEG studies.

## Competing Interests

The author declares no competing interests.

## Funding

This research received no specific grant from any funding agency, public, commercial, or not-for-profit.

## Data availability

MPI–LEMON EEG data are publicly available (see original citation).

## Supplementary Materials

### S1. LEMON EEG ingestion and time–frequency scalp topographies

#### Source data

We analyzed the consortium-provided *preprocessed* resting EEG from the Leipzig Mind–Brain–Body (MPI–LEMON) project (Babayan and et al., 2019), obtained via the GWDG mirror^1^ (`EEG_Preprocessed_BIDS_ID/EEG_Preprocessed`). Eyes-open (EO) and eyes-closed (EC) recordings were distributed as EEGLAB `.set/.fdt` pairs. Only participants with both EO and EC files were included.

#### Loading and referencing

Data were read with MNE–Python’s EEGLAB reader (Delorme and Makeig, 2004). We retained EEG channels only, applied the standard 10–10 montage (`standard_1005` with `on_missing=‘ignore’`), and re-referenced to the common average of retained EEG sensors. We relied on the consortium’s preprocessing (downsampling, band-pass filtering, channel cleaning, and ICA-based ocular/ECG removal per Babayan and et al., 2019); no additional artifact removal was performed. All subsequent analyses used these continuous preprocessed recordings.

LEMON’s EO/EC recordings consist of sixteen 60 s blocks (8 EO, 8 EC) recorded in alternation. In the preprocessed files, EO blocks are concatenated into `sub-*_EO.set` and EC blocks into `sub-*_EC.set`; EEGLAB inserts `boundary` events at each join. To avoid edge transients in time–frequency estimates, each condition was segmented into non-overlapping 10 s windows, and any window overlapping a `boundary` annotation was discarded.

#### Canonical channel mask

To ensure comparability across subjects and methods, we enforced a fixed 59-channel canonical layout. For each subject, we intersected this list with available channels after EO/EC pairing and constructed a Boolean mask; no spatial interpolation was performed. Both the dictionary rows (leadfield-projected LB basis in the sensor space) and the observed topographies were subset and ordered according to this per-subject mask, yielding identical channel order across methods (LB, SPH, PCA, ICA) and conditions.

#### Morlet time–frequency decomposition

For each condition (EO/EC), each surviving 10 s window was transformed with a complex Morlet wavelet using `mne.time_frequency.tfr_array_morlet` (single precision, `n_jobs=1`). We used a logarithmically spaced frequency vector *f* ∈ [2, 30] Hz with *F* =15 points. To balance temporal and spectral resolution across the band, a constant-*Q* schedule was used with frequency-proportional cycles bounded between 3 and 10, i.e., *n*_cycles_(*f*) = min {max(*f/*2, 3), 10} (Torrence and Compo, 1998; Cohen, 2019). Time–frequency power was averaged over time within each 10 s window to form one topography of length-*M* per analyzed frequency; these *F* topographies were then concatenated across frequencies, yielding a raw topography matrix *Y* ∈ ℝ^(*T F*)×*M*^ per subject, where *T* is the number of kept windows and *M* the subject’s kept channels. In addition, we also produced band-aggregated topographies by averaging power across canonical bands (*δ* 2–4 Hz; *θ* 4–7 Hz; *α* 8–12 Hz; *β* 13–30 Hz), producing a (*T* × 4) × *M* matrix per condition. No sensor-space baseline (dB) conversion was required for downstream span analyses because all projections operate on row-wise *z*-scored topographies (centering and scaling across channels per row, where each row indicates a specific timefrequency pair to remove an offset across channels) to emphasize spatial pattern rather than absolute power.

A subject-level manifest (.csv) recorded the number of retained channels, epochs per condition, frequency grid, cycle rule, decimation factor, and file paths. Across all eligible preprocessed LEMON participants, *N* =202 contributed at least one valid EO and one valid EC window and were retained for group analysis.

#### Reproducibility

The loading and TFR extraction pipeline is implemented in `lemon_batch_tfr.py` (Python 3; `mne, numpy, scipy, pandas`). Default arguments were `--win-len 10, --fmin 2, --fmax 30, --n-freqs 15, --cycles-rule constQ`, and `--decim 2`. Boundary-aware epoching and canonical channel masking were enabled by default. Scripts and channel lists are available in the project repository.

#### Montage harmonization

All analyses are performed in a harmonized 10–10 coordinate system so that the forward-model dictionary, observed topographies, and comparator bases (LB, SPH, PCA, ICA) operate on the same channels in the same order.

#### Canonical 59-channel set

We define a canonical scalp set of *M* =59 EEG channels by: (i) starting from the `standard_1005` montage; (ii) excluding non-EEG sensors (e.g., EOG) and site-specific extras; and (iii) retaining only 10–10 positions that are consistently present across the cohort with stable 3D positions under `standard_1005`.^2^ This set provides near-uniform coverage of frontal, central, parietal, temporal, and occipital regions and closely matches common 64-channel layouts while omitting ocular/site-specific sensors that could confound representational analyses.

#### Subject-specific masking

For each subject, we construct a Boolean presence mask over the canonical 59 channels and apply the same mask to both the dictionary rows and the observed data columns. This yields a subject-specific *m*_used_ ≤ 59 (to accommodate rare missing/bad channels) while maintaining perfect row/column alignment across methods. The forward operator *L* and the LB dictionary *D* = *L*Φ are built in the canonical channel order; all comparators (SPH/PCA/ICA) are evaluated on the identical reduced set per subject to ensure fair comparisons.

#### Downsampled montages

To assess robustness to spatial sampling density, we also analyze standard downsampled subsets: a 32-channel set (`canonical_32.txt`, 10–10 “BioSemi-32”-style)^3^ and a 19-channel set (`canonical_19.txt`, international 10–20)^4^. For these, we either restrict fixed dictionaries (LB/SPH) to the kept rows or (for PCA/ICA) recompute the basis on the reduced montage, always reusing the same subject-specific mask so that every method sees exactly the same channels. This harmonized design respects known spatial-sampling limits of EEG (e.g., Nunez and Srinivasan, 2006; Srinivasan et al., 1998) and maximizes cross-subject comparability and reproducibility.

### S2. Laplace–Beltrami basis, whole-cortex modes, and EEG sensor dictionary

We build a geometry-aligned basis once on the *fsaverage* template and obtain a *sensor-space, EEG-specific* dictionary by forward projection through an MNE/BEM head model. The complete procedure is summarized in **Algorithm 1** (see “Algorithm 1. Template LB eigenbasis → whole-cortex modes → sensor dictionary”).

#### Cortical surface and discretization

We computed cortical harmonics on the *fsaverage* template (left/right pial meshes; FreeSurfer) (Fischl, 2012; Dale et al., 1999), treating each hemisphere as a two-dimensional Riemannian manifold embedded in ℝ^3^. Let Δ_ℳ_ denote the Laplace–Beltrami operator on the cortical surface ℳ. The continuous eigenproblem

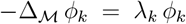

was discretized with a standard finite-element (FEM) cotangent scheme on the triangle mesh, yielding the generalized symmetric eigenproblem

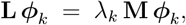

where **L** is the cotangent stiffness matrix and **M** the (consistent) mass matrix (Meyer et al., 2003; Reuter et al., 2006, 2009). We solved for the lowest non-trivial eigenpairs per hemisphere, discarded the constant (DC) mode, and mass-normalized eigenvectors so that 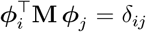. In our implementation, we retained *K* = 60 modes per hemisphere (coarse–to–mid spatial frequencies on cortex).

#### Whole-cortex mode assembly

To obtain bilaterally consistent modes with a single index *k* across the hemispheres, we pair LH/RH eigenmodes by eigenvalue order (*k* ↔ *k*) and form whole-cortex combinations after reflecting RH across the mid-sagittal plane and matching vertices by Euclidean nearest neighbor. Let *T*_*R*→*L*_ = NN_*L*_ ◦Mirror, denote the alignment operator (“mirror then nearest-neighbor onto the opposite hemisphere”; from RH to LH; see “*Implementation detail”* below). Writing 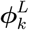 and 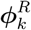 for the *k*-th LH/RH modes, we define a global sign

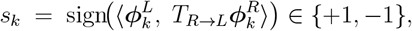

and concatenate

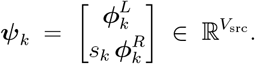

Reflection+nearest-neighbor “locks” the arbitrary orientation of hemispheric pairs, producing bilaterally consistent, anatomy-aligned modes that preserve the low-to-high spatial-frequency ordering (Pang et al., 2023).

#### Implementation detail

For each mode index *k*, we align hemispheres in 3D source coordinates by reflecting RH vertices across the mid–sagittal plane and, for each LH vertex *i* with coordinate 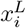, selecting the vertex index *j*^∗^(*i*), such that 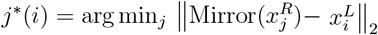. We then evaluate 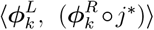 over used vertices and set *s*_*k*_ accordingly. The nearest-neighbor map is used only to compute the per-mode sign *s*_*k*_ reliably; the final whole-cortex column is formed by stacking LH and the sign-aligned RH, without any hemispheric averaging. Reflecting across the mid–sagittal plane followed by nearestneighbor search in source coordinates reliably aligns homologous banks of gyri/sulci at negligible cost.

#### Alignment to MNE source space and forward projection (dictionary formation)

MNE– Python was used to set up a cortical *source space* on *fsaverage* with `spacing=ico4` (about ∼ 2500 vertices/hemisphere) (Gramfort et al., 2014). We reindex ***ψ***_*k*_ to match the exact vertex order of the MNE source grid, resulting in the matrix 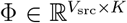 (columns are whole-cortex, sign-aligned modes). The forward (leadfield) model for EEG was computed with a three-layer BEM (conductivities [0.3, 0.006, 0.3] S*/*m for scalp, skull, brain) and fixed (surface-normal) orientation, enforcing a minimum 5 mm distance from the inner skull (Vorwerk et al., 2014)^5^.Let. 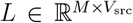 be the resulting EEG leadfield (rows: sensors in canonical order; columns: source vertices), and 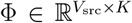 collect the reindexed whole-cortex LB modes. The *sensor-space LB dictionary* is then

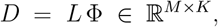

whose *k*-th column is the scalp topography generated by the *k*-th cortical harmonic after realistic volume conduction. In analysis, we projected observed topographies by ordinary least squares onto an orthonormalized version of *D* (thin SVD with numerical rank truncation at 10^−8^ × *σ*_max_), preserving LB ordering while ensuring well-conditioned span tests.

Pang et al. (Pang et al., 2023) computed eigenmodes on a high-resolution cortical template (e.g., fs LR 32k) and analyzed cortical/fMRI patterns in cortical coordinates; symmetric/anti-symmetric combinations are discussed, but the focus was on cortex-space representational and dynamical models. Here, we (i) compute LB modes on *fsaverage* (FEM cotangent discretization; mass normalization), (ii) explicitly *forward-project* them through a realistic EEG head model to obtain a *sensor-space* dictionary *D*, and (iii) evaluate representational efficiency and reliability by OLS span tests at the *scalp*.

In practice, the leading ∼ 10–20 harmonics capture scalp EEG’s resolvable spatial bandwidth; higher modes encode detail strongly attenuated by the head volume conductor and montage sampling (Srinivasan et al., 1998; Iivanainen et al., 2021a; Pang et al., 2023).

#### Reproducibility

All steps follow standard discrete LB formulations on cortical meshes with mass normalization (Meyer et al., 2003; Reuter et al., 2006, 2009) and established MNE/BEM procedures for EEG forward modeling on *fsaverage* (Gramfort et al., 2014; Vorwerk et al., 2014). The entire procedure is implemented in two self-contained scripts: `compute _phi_lapy.py` (LB eigenmodes on *fsaverage*, hemisphere pairing via mirror+nearest neighbor, global sign alignment, reindexing to MNE source space) and `make_fsaverage_lb_dictionary.py` (forward model and *D* = *L*Φ construction).

Our goal is a *sensor–space, EEG-specific, fsaverage-based* dictionary that plugs directly into an MNE–Python head model and is usable without subject MRIs at test time. This requires three design choices not covered by the BrainEigenmodes reference code of Pang et al. (2023):

- **Mesh and toolchain**. We operate on *FreeSurfer fsaverage* (LH/RH pial) and an MNE `ico4` source space for EEG forward modeling (Gramfort et al., 2014). We implement a pure-Python FEM LB solver (via `lapy+scipy`), avoiding external toolchains (e.g., Workbench/MATLAB) and guaranteeing exact vertex indexing consistent with MNE’s source space, which is crucial for constructing *D* = *L*Φ and for montage-aware masking.
- **Whole-cortex stacked assembly in source order**. For EEG we require columns that already span both hemispheres in the exact MNE vertex order. We therefore build whole-cortex stacked combinations explicitly with mirror+NN alignment (operators *T*_*R*→*L*_) and reindex directly to the MNE grid prior to forward projection.
- **Forward projection**. After assembling Φ on the MNE grid, we explicitly propagate modes through a three-layer EEG BEM to form *D* = *L*Φ (Gramfort et al., 2014; Vorwerk et al., 2014). We then orthonormalize *D* in sensor space with a strict relative cutoff to obtain a montage-specific, well-conditioned basis for span tests and coefficient reliability.

##### Algorithm 1.

Template LB eigenbasis → whole-cortex modes → sensor dictionary

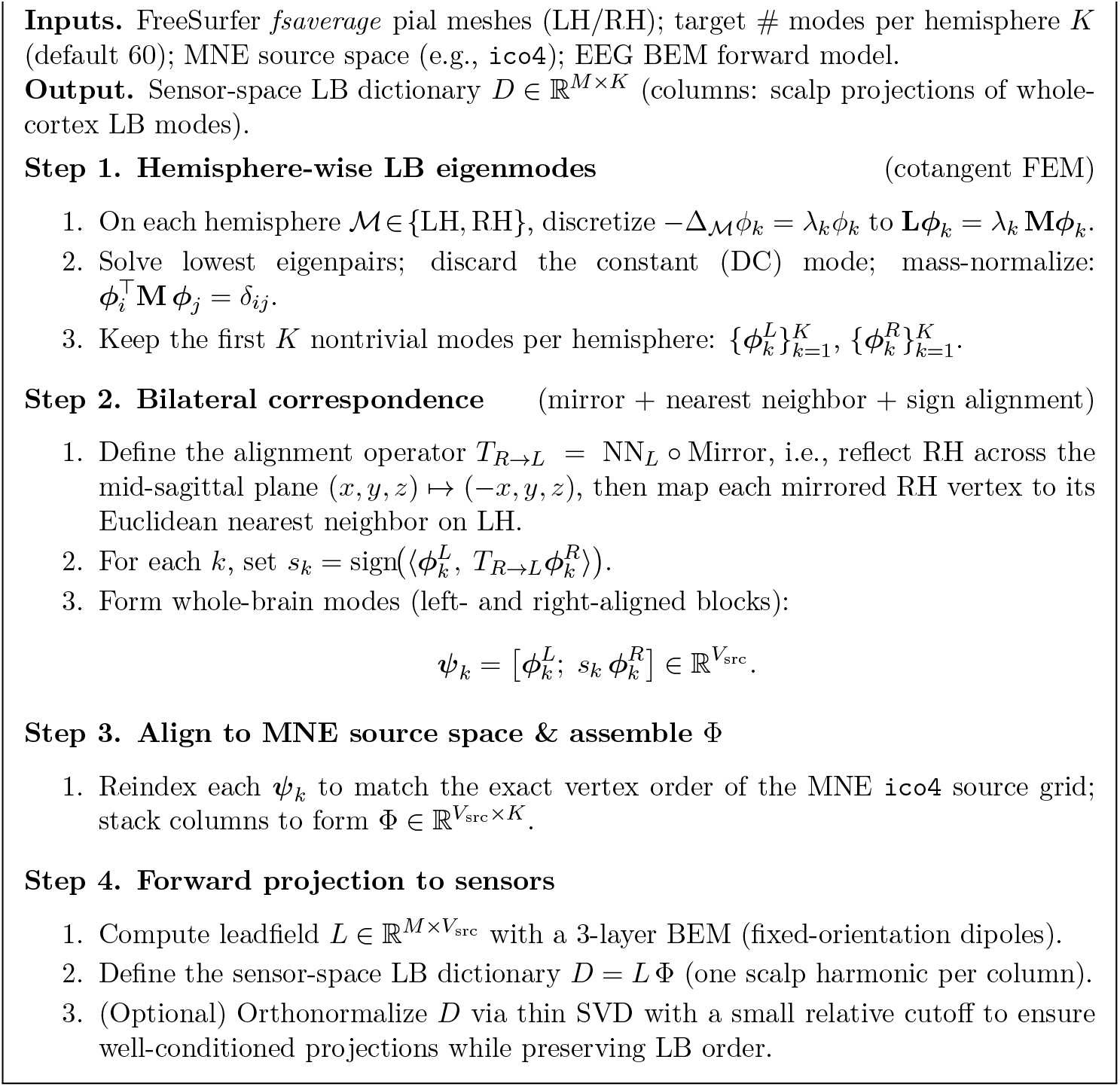

**Table S1:** LB EO–EC reliability per mode (59-ch): ICC(3,1) with percentile bootstrap 95% CIs for *α, β, δ, θ* bands. Modes 1–20 are shown here (native LB order).

| Mode | $\alpha$ | | $\beta$ | | $\delta$ | | $\theta$ | |
| --- | --- | --- | --- | --- | --- | --- | --- | --- |
| 1 | 0.472 | [0.361, 0.577] | 0.479 | [0.360, 0.582] | 0.453 | [0.338, 0.555] | 0.454 | [0.349, 0.578] |
| 2 | 0.430 | [0.304, 0.538] | 0.445 | [0.312, 0.556] | 0.405 | [0.268, 0.511] | 0.431 | [0.283, 0.540] |
| 3 | 0.366 | [0.208, 0.521] | 0.368 | [0.201, 0.513] | 0.341 | [0.184, 0.474] | 0.366 | [0.206, 0.505] |
| 4 | 0.560 | [0.436, 0.663] | 0.512 | [0.378, 0.630] | 0.557 | [0.442, 0.662] | 0.578 | [0.466, 0.677] |
| 5 | 0.553 | [0.401, 0.671] | 0.539 | [0.375, 0.657] | 0.527 | [0.380, 0.650] | 0.548 | [0.399, 0.668] |
| 6 | 0.439 | [0.288, 0.567] | 0.472 | [0.356, 0.578] | 0.448 | [0.327, 0.556] | 0.473 | [0.341, 0.588] |
| 7 | 0.456 | [0.286, 0.609] | 0.473 | [0.313, 0.617] | 0.450 | [0.290, 0.605] | 0.466 | [0.277, 0.609] |
| 8 | 0.607 | [0.508, 0.696] | 0.617 | [0.515, 0.701] | 0.594 | [0.489, 0.693] | 0.624 | [0.526, 0.715] |
| 9 | 0.313 | [0.101, 0.484] | 0.316 | [0.111, 0.490] | 0.294 | [0.091, 0.475] | 0.314 | [0.093, 0.497] |
| 10 | 0.594 | [0.478, 0.687] | 0.566 | [0.430, 0.673] | 0.615 | [0.498, 0.712] | 0.622 | [0.503, 0.711] |
| 11 | 0.342 | [0.194, 0.478] | 0.366 | [0.231, 0.484] | 0.366 | [0.236, 0.476] | 0.372 | [0.237, 0.487] |
| 12 | 0.466 | [0.335, 0.593] | 0.478 | [0.345, 0.592] | 0.447 | [0.317, 0.575] | 0.462 | [0.317, 0.600] |
| 13 | 0.512 | [0.390, 0.622] | 0.494 | [0.370, 0.598] | 0.502 | [0.382, 0.607] | 0.513 | [0.400, 0.623] |
| 14 | 0.326 | [0.143, 0.486] | 0.374 | [0.202, 0.523] | 0.322 | [0.149, 0.492] | 0.320 | [0.145, 0.482] |
| 15 | 0.452 | [0.344, 0.558] | 0.426 | [0.315, 0.532] | 0.445 | [0.338, 0.546] | 0.462 | [0.350, 0.564] |
| 16 | 0.572 | [0.416, 0.696] | 0.586 | [0.427, 0.703] | 0.574 | [0.436, 0.693] | 0.561 | [0.416, 0.682] |
| 17 | 0.561 | [0.435, 0.669] | 0.541 | [0.432, 0.653] | 0.570 | [0.439, 0.691] | 0.566 | [0.445, 0.680] |
| 18 | 0.381 | [0.231, 0.525] | 0.389 | [0.248, 0.526] | 0.383 | [0.225, 0.526] | 0.383 | [0.229, 0.537] |
| 19 | 0.483 | [0.331, 0.624] | 0.467 | [0.308, 0.619] | 0.455 | [0.299, 0.588] | 0.470 | [0.311, 0.611] |
| 20 | 0.555 | [0.401, 0.687] | 0.541 | [0.380, 0.672] | 0.570 | [0.416, 0.687] | 0.572 | [0.409, 0.689] |

### S3. ICC(3,1)

To assess EO/EC test–retest of mode × band coefficients, we use ICC(3,1) (two-way mixed, single measurement, *consistency*) (Shrout and Fleiss, 1979), treating EO and EC as the two “raters.” For each subject *s*, mode *k*, and band *b*, we form *band-mean* OLS coefficients by averaging *c*_*s,i,k*_(*t, f*) over all windows *t* and all frequencies *f* in band *b*, separately for EO (*i* = EO) and EC (*i* = EC). ICC is then computed *across subjects* on these paired EO/EC values for each (*k, b*).

Let *x*_*is*_ denote the coefficient for rater *i* ∈ {EO, EC} and subject *s* = 1, … , *N* , with *q*=2 “raters.” In the two-way mixed ANOVA, *MS*_subj_ and *MS*_err_ are the mean squares for subjects and residual error, respectively, and

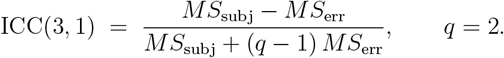

(With 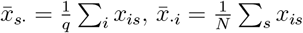 ,and 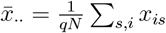

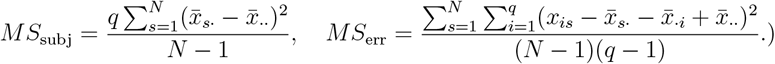

### S4. Bootstrap procedure (subject-level)

All confidence intervals (CIs) are obtained via a nonparametric percentile bootstrap that resamples subjects with replacement, preserving within-subject dependence (all windows, frequencies/bands, and EO/EC for a subject travel together). For each montage and method, and for each condition *c* ∈ {EO, EC}:

**Table S2:** SPH EO–EC reliability per mode (59-ch): ICC(3,1) with percentile bootstrap 95% CIs for *α, β, δ, θ* bands. Rows list SPH modes 1–20 in degree-major order (covering degrees *ℓ*=0–4).

| Mode | $\alpha$ | | $\beta$ | | $\delta$ | | $\theta$ | |
| --- | --- | --- | --- | --- | --- | --- | --- | --- |
| 1 | 0.455 | [0.340, 0.566] | 0.523 | [0.404, 0.620] | 0.489 | [0.372, 0.589] | 0.441 | [0.307, 0.565] |
| 2 | 0.485 | [0.332, 0.613] | 0.455 | [0.312, 0.596] | 0.479 | [0.338, 0.600] | 0.479 | [0.341, 0.599] |
| 3 | 0.504 | [0.358, 0.616] | 0.546 | [0.420, 0.662] | 0.516 | [0.387, 0.618] | 0.513 | [0.389, 0.616] |
| 4 | 0.458 | [0.323, 0.579] | 0.473 | [0.326, 0.598] | 0.434 | [0.317, 0.556] | 0.436 | [0.303, 0.560] |
| 5 | 0.432 | [0.223, 0.580] | 0.445 | [0.265, 0.576] | 0.446 | [0.231, 0.612] | 0.465 | [0.258, 0.617] |
| 6 | 0.598 | [0.473, 0.703] | 0.591 | [0.462, 0.710] | 0.575 | [0.441, 0.687] | 0.609 | [0.473, 0.723] |
| 7 | 0.535 | [0.406, 0.659] | 0.510 | [0.380, 0.639] | 0.558 | [0.429, 0.676] | 0.548 | [0.427, 0.656] |
| 8 | 0.463 | [0.342, 0.574] | 0.461 | [0.305, 0.600] | 0.442 | [0.326, 0.553] | 0.442 | [0.314, 0.552] |
| 9 | 0.415 | [0.276, 0.565] | 0.380 | [0.240, 0.527] | 0.381 | [0.211, 0.555] | 0.400 | [0.262, 0.554] |
| 10 | 0.421 | [0.277, 0.544] | 0.401 | [0.265, 0.527] | 0.416 | [0.279, 0.539] | 0.411 | [0.268, 0.543] |
| 11 | 0.430 | [0.236, 0.591] | 0.421 | [0.223, 0.575] | 0.426 | [0.206, 0.596] | 0.456 | [0.248, 0.628] |
| 12 | 0.474 | [0.313, 0.618] | 0.447 | [0.284, 0.591] | 0.476 | [0.327, 0.602] | 0.479 | [0.329, 0.612] |
| 13 | 0.581 | [0.430, 0.699] | 0.611 | [0.481, 0.717] | 0.579 | [0.452, 0.685] | 0.580 | [0.449, 0.700] |
| 14 | 0.481 | [0.353, 0.591] | 0.485 | [0.332, 0.615] | 0.455 | [0.330, 0.567] | 0.453 | [0.336, 0.569] |
| 15 | 0.379 | [0.216, 0.521] | 0.392 | [0.226, 0.533] | 0.369 | [0.209, 0.510] | 0.363 | [0.183, 0.505] |
| 16 | 0.535 | [0.424, 0.643] | 0.537 | [0.437, 0.639] | 0.516 | [0.398, 0.631] | 0.524 | [0.412, 0.644] |
| 17 | 0.486 | [0.299, 0.647] | 0.560 | [0.351, 0.719] | 0.466 | [0.261, 0.627] | 0.494 | [0.318, 0.648] |
| 18 | 0.272 | [0.145, 0.408] | 0.269 | [0.141, 0.399] | 0.269 | [0.136, 0.407] | 0.272 | [0.140, 0.417] |
| 19 | 0.422 | [0.231, 0.568] | 0.432 | [0.246, 0.570] | 0.425 | [0.221, 0.578] | 0.453 | [0.265, 0.601] |
| 20 | 0.565 | [0.442, 0.674] | 0.535 | [0.414, 0.653] | 0.517 | [0.379, 0.622] | 0.554 | [0.419, 0.656] |

#### (i) Subject-level statistics

For efficiency curves, compute the subject-level mean 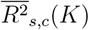 by averaging row-wise *R*^2^(*K*; *y*) over that subject’s retained rows (windows × frequencies or bands). Define 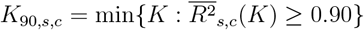 with right-censoring at *K* = *r* (the SVD effective rank of the orthonormalized basis for that subject). For reliability, assemble the paired EO/EC band-mean coefficients across subjects and compute ICC(3,1) per mode × band.

#### (ii) Resampling

For *b* = 1, … , *B* with *B* = 2000, draw *N* subject IDs with replacement to form a bootstrap cohort *S*_*b*_ (EO and EC always kept paired). Recompute group summaries on *S*_*b*_:

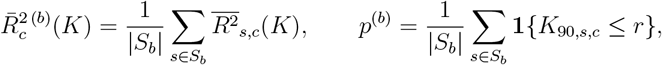

and the mean (and normalized) *K*_90_ among attainers in *S*_*b*_. For reliability, recompute ICC(3,1) per mode × band on *S*_*b*_.

#### (iii) Confidence intervals

Report 95% CIs using the percentile bootstrap (2.5th/97.5th percentiles across 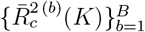 at each *K*, across {*p*^(*b*)^}, across the bootstrap distribution of mean *K*_90_ and *K*_90_*/M* among attainers, and across ICCs). Pointwise CIs in *K* are reported for *R*^2^(*K*).

### S5. Implementation code information

#### Environment and versions

Python 3.13.5 with NumPy, SciPy, pandas, scikit–learn, and MNE–Python. We used `mne.datasets.fetch_fsaverage` to install `fsaverage`.

**Table S3:**
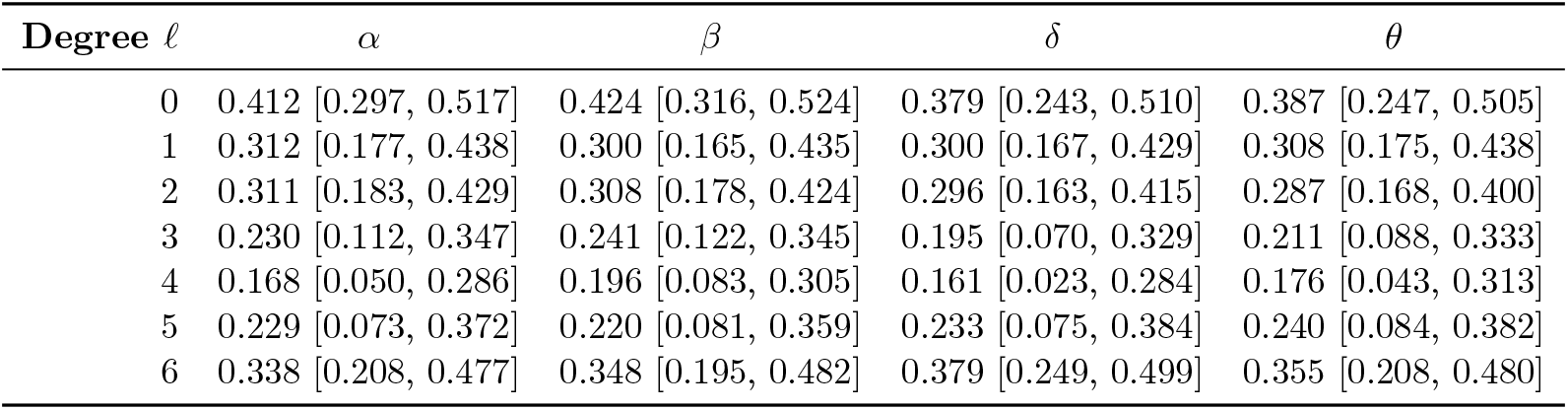
Degree-aggregated SPH reliability (59-ch): ICC(3,1) with 95% bootstrap CIs by spherical degree *ℓ* and band. Aggregation uses 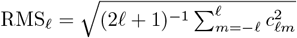. Degree *ℓ*=7 is omitted because, at 59 channels, it is sampling-limited and collapses to ICC= 0 with a zero-width CI.

**Table S4:** Projection throughput (59-ch) at *K*=20: median seconds per 1,000 rows for basis formation (thin SVD/QR) of the basis *B* plus OLS projection *Y* → (*Y Q*)*Q* , single-threaded BLAS, averaged over 5 repeats and summarized as the subject-level median (*N* =202). The results indicate that LB projections are lightweight; ICA is an order of magnitude slower due to component fitting.

| Method | EC (sec/1,000 rows) | EO (sec/1,000 rows) |
| --- | --- | --- |
| SPH | 0.003396 | 0.003433 |
| LB | 0.001476 | 0.001497 |
| PCA | 0.004933 | 0.004898 |
| ICA | 0.100564 | 0.108917 |

#### Lead-field and source space

Three-layer BEM (scalp, skull, brain; conductivities 0.3, 0.006, 0.3 S*/*m), `fsaverage` source space at `ico4` (∼ 2,500 vertices/hemisphere). EEG forward model with fixed–orientation dipoles aligned to the surface normal and minimum 5 mm source–to–inner–skull distance:

- `mne.make_bem_model(ico=4, conductivity=(0.3,0.006,0.3))` ,
- `mne.make_forward_solution(info, trans=‘fsaverage’, src, bem, eeg=True, meg=False`
- `mne.convert_forward_solution(…, surf_ori=True, force_fixed=True)` .

#### LB eigenmodes and sensor dictionary

- `compute_phi_lapy.py`: computes cortical Laplace-Beltrami eigenmodes on *fsaverage* using a cotangent FEM (LaPy), per hemisphere, drops the DC mode, maps to the MNE source-space vertex order, and forms 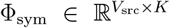 as stacked whole-cortex columns by pairing LH/RH modes *k* and sign-aligning RH to LH via mirror+nearest–neighbor on the source grid.
- `make_fsaverage_lb_dictionary.py`: builds the sensor dictionary *D* = *L*Φ_sym_ (rows reindexed to the canonical channel order) and writes . `npz` files: `fsaverage_phi_sym_K{K}.npz` (Φ, eigenvalues) and `fsaverage_D_sym_K{K}_M{M}.npz` (*D*, channels, metadata).

#### LEMON preprocessing and topographies

- `lemon_batch_tfr.py`: downloads preprocessed EEG from the public GWDG mirror, performs boundary–aware epoching of EO/EC resting runs (default 10 s non-overlapping windows; epochs overlapping “boundary” annotations are dropped), computes Morlet TFR power with log-spaced *f* ∈ [2, 30] Hz and constQ cycles (*n*_cycles_(*f*) = clip(*f/*2, 3, 10)), and optionally aggregates bands (*δ* : [2, 4], *θ* : [4, 7], *α* : [8, 12], *β* : [13, 30] Hz).
- Output per subject: `sub-_Y_tfr.npz` containing raw–frequency and band-averaged topographies for EO/EC, the canonical channel list (`canonical_channels`), and a per-subject boolean `subject_mask` onto the canonical montage.

#### OLS span tests and group summaries

- `run_lb_fits.py`: LB representational efficiency. Projects onto the span of the *leading K* LB columns in *native order*, re-orthonormalized at each *K* via no-pivot QR (i.e., “take the first *K* columns, QR, and project onto that *K*–dimensional span”). Computes the full *R*^2^(*K*) curve per subject/condition, thresholds *K*_70*/*90_ from the full curve, and stores LB values at SPH degree checkpoints *K* ∈ {1, 4, 9, 16, 25, 36, 49, … }.
- `run_baselines_pca_ica_sph.py`: PCA/ICA/SPH span tests with identical masking and montages. PCA uses a thin SVD of column-centered *Y* (then QR-cleanup); ICA uses FastICA (unit-variance whitening) then QR on the mixing topographies; SPH builds a real spherical-harmonics design in degree-major order and applies no–pivot QR. PCA/ICA thresholds *K*_70*/*90_ come from the full curve; SPH *R*^2^(*K*) is recorded at degree–block checkpoints.
- `bootstrap_curves.py`: merges LB & baseline comparators and computes group-mean *R*^2^(*K*) with nonparametric bootstrap CIs (subject-level resampling), attainment rates, and censored summaries for *K*_70*/*90_. Writes `group_curves_bootstrap.csv` and `attainment_k90_bootstrap.csv`.

#### Shared-basis coefficients and reliability (ICC)

- `run_shared_coeffs_icc.py`: computes EO/EC coefficients in the shared fixed coordinates for LB and SPH, forms band–mean coefficient vectors, and computes ICC(3,1) per band × mode with bootstrap CIs. For SPH it reports all modes (and also tags each mode with its spherical degree and whether it is a degree-block endpoint).
- `sph_degree_icc.py`: rotation-invariant SPH summary. Aggregates within each spherical degree by the root-mean-squared (RMS) coefficients across the 2*ℓ* + 1 modes, then computes EO/EC ICC(3,1) per degree with bootstrap CIs.

#### Computational efficiency benchmarking

- `benchmark_methods.py`: benchmarks LB/PCA/ICA/SPH “preparation+projection” time per 1,000 rows (median across subjects), with options for `--rows, --repeat`, and montage subsets. LB/SPH use native-order no-pivot QR; PCA uses SVD; ICA uses FastICA→QR.

#### Canonical montages and subsets

We provide `canonical_59.txt, canonical_32.txt`, and `canonical_19.txt` (one channel name per line). All scripts accept `--subset-file` to apply an additional mask in the canonical dictionary order (`ALL59, canonical 32, canonical 19` in outputs).

## Footnotes

1 https://ftp.gwdg.de/pub/misc/MPI-Leipzig_Mind-Brain-Body-LEMON/EEG_MPILMBB_LEMON/

2 The exact list is provided in canonical 59.txt and reproduced here: Fp1, Fp2, F7, F3, Fz, F4, F8, FC5, FC1, FC2, FC6, C3, Cz, C4, T8, CP5, CP1, CP2, CP6, AFz, P7, P3, Pz, P4, P8, PO9, O1, Oz, O2, PO10, AF7, AF3, AF4, AF8, F5, F1, F2, F6, FT7, FC3, FC4, FT8, C5, C1, C2, C6, CP3, CPz, CP4, TP8, P5, P1, P2, P6, PO7, PO3, POz, PO4, PO8.

3 The exact list is provided in `canonical_32.txt` and reproduced here: Fp1, Fp2, AF3, AF4, F7, F3, Fz, F4, F8, FC5, FC1, FC2, FC6, C3, Cz, C4, T8, CP5, CP1, CP2, CP6, P7, P3, Pz, P4, P8, PO7, PO3, POz, PO4, O1, O2.

4 The exact list is provided in `canonical_19.txt` and reproduced here: Fp1, Fp2, F7, F3, Fz, F4, F8, FT7, C3, Cz, C4, T8, P7, P3, Pz, P4, P8, O1, O2.

5 In code: we built the BEM with `mne_make_bem_model(ico=4, conductivity=(0.3,0.006,0.3)) , obtained the forward with mne.make_forward_solution(info, trans=‘fsaverage’, src, bem, eeg=True, meg=False, mindist=5.0`), converted to fixed surface-normal orientation via `mne.convert_forward_solution(…, surf ori=True, force_fixed=True`), extracted *L* as fwd[‘sol’][‘data’], and finally formed *D* = *L*Φ.

## Notes

### Competing Interest Statement

The authors have declared no competing interest.

